# Using a supervised principal components analysis for variable selection in high-dimensional datasets reduces false discovery rates

**DOI:** 10.1101/2020.05.15.097774

**Authors:** Insha Ullah, Kerrie Mengersen, Anthony Pettitt, Benoit Liquet

## Abstract

High-dimensional datasets, where the number of variables ‘*p*’ is much larger compared to the number of samples ‘*n*’, are ubiquitous and often render standard classification and regression techniques unreliable due to overfitting. An important research problem is feature selection — ranking of candidate variables based on their relevance to the outcome variable and retaining those that satisfy a chosen criterion. In this article, we propose a computationally efficient variable selection method based on principal component analysis. The method is very simple, accessible, and suitable for the analysis of high-dimensional datasets. It allows to correct for population structure in genome-wide association studies (GWAS) which otherwise would induce spurious associations and is less likely to overfit. We expect our method to accurately identify important features but at the same time reduce the False Discovery Rate (FDR) (the expected proportion of erroneously rejected null hypotheses) through accounting for the correlation between variables and through de-noising data in the training phase, which also make it robust to outliers in the training data. Being almost as fast as univariate filters, our method allows for valid statistical inference. The ability to make such inferences sets this method apart from most of the current multivariate statistical tools designed for today’s high-dimensional data. We demonstrate the superior performance of our method through extensive simulations. A semi-real gene-expression dataset, a challenging childhood acute lymphoblastic leukemia (CALL) gene expression study, and a GWAS that attempts to identify single-nucleotide polymorphisms (SNPs) associated with the rice grain length further demonstrate the usefulness of our method in genomic applications.

**Author summary:** An integral part of modern statistical research is feature selection, which has claimed various scientific discoveries, especially in the emerging genomics applications such as gene expression and proteomics studies, where data has thousands or tens of thousands of features but a limited number of samples. However, in practice, due to unavailability of suitable multivariate methods, researchers often resort to univariate filters when it comes to deal with a large number of variables. These univariate filters do not take into account the dependencies between variables because they independently assess variables one-by-one. This leads to loss of information, loss of statistical power (the probability of correctly rejecting the null hypothesis) and potentially biased estimates. In our paper, we propose a new variable selection method. Being computationally efficient, our method allows for valid inference. The ability to make such inferences sets this method apart from most of the current multivariate statistical tools designed for today’s high-dimensional data.

## Introduction

Many new challenges have been raised over recent years by high-dimensional data, especially so-called ‘large *p* small *n*’ data with thousands or tens of thousands of features but a limited number of samples. This brings the standard analysis tools to a halt or deteriorates the learning process as the learning algorithm becomes prone to over-fitting due to the abundance of irrelevant and redundant features. This means that the good performance of a fitted model could be limited to the training data set and its performance could be very poor for new unseen samples. An obvious challenge in such applications is feature selection, which is designed to filter out the features that have little to no correlation with the response variable based on the available data. Feature selection is an integral component of modern statistical research, and methods that are computationally less demanding and that obtain reliable results are urgently needed.

Recently, considerable effort has been directed to identifying important variables that contribute useful information towards a response variable of interest. The approaches that are commonly used can be broadly divided into two categories: univariate filters and multivariate feature selection methods. The univariate filters independently assess each feature based on its relevance to the response and reduce the dimension by filtering out all those features that are not strongly associated with a response variable [for example, see 1, 2, 3]. Owing to their simplicity, interpretability and computational efficiency, the filters are more commonly used in the analysis of high-dimensional datasets. One of the drawbacks of univariate filters, however, is that dependencies among variables are ignored because the variables are assessed one at a time. As a result, a substantial amount of information contained in the data is left under utilized. Moreover, ignoring correlations between variables may lead to biased estimates and incorrect or underpowered inferences.

Multivariate variable selection methods overcome this drawback and use all the variables simultaneously to choose the best subset of variables. Many multivariate variable selection methods have been proposed. For example, the lasso regression [4] and its improved variants like the elastic-net regression [5] are widely used because they produce sparse solutions and perform the model selection simultaneously with parameter estimation. Random forest models, which use an ensemble of classification or regression trees, are also popular [6]. The algorithm returns measures of variable importance and has been used in many applications, including gene selection [7]. However, both lasso regression and random forest were designed for datasets of a moderate size (hundreds of variables) and rely on computationally cumbersome methods such as cross-validation for tuning penalty parameters. For high-dimensional datasets, these methods may be statistically inaccurate [8, 9].

An intermediate approach reduces the dimensionality of data to a manageable size by first using a univariate filter, which is then followed by an appropriate multivariate method to perform the final selection. This is adopted by [10] who introduced iteratively sure independent screening (ISIS) in the context of linear regression models and showed that it is a suitable variable selection strategy for high-dimensional settings. The procedure couples an independent screening of a large number of variables based on their marginal correlations with the final variable selection and parameter estimation using a more sophisticated multivariate technique such as elastic-net logistic regression [5]. The method has been used for variable selection with a range of high-dimensional datasets [for example, see 11]. The ISIS idea is also extended to logistic regression of case control data [8]. Other similar screening techniques to accelerate the computation at the second stage of optimization for final variable selection and parameters estimation include SAFE rules [12] and strong rules [13]. However, these methods still suffer from the drawbacks of computational inefficiency and statistical inaccuracy at the second stage of the final variable selection if the number of filtered variables is large compared to the number of available samples.

Principal component analysis (PCA) has been used for dimension reduction in high-dimensional data because it is statistically coherent, computationally fast and scalable [14]. A disadvantage of using PCA as a dimension reduction technique is that the principal components (PCs) are linear combinations of all original variables and often may not be easy to interpret. Even if the PCs are interpretable, which is sometimes the case for the first few PCs, one often needs interpretation in terms of original variables. For example, one might be interested in identifying genes that are associated with a particular disease. Attempts have been made to use PCA for variable selection rather than variable extraction [for example, see 15, 16]. However, these methods are computationally infeasible for high-dimensional datasets. Shen and Huang [17] introduced sparse PCA that produce PCs with sparse loadings (a.k.a coefficients) through forcing near zero loading to be exactly zero. The variables thus associated with the non-zero loadings in the first few PCs are considered informative. More recently, [18] have adapted PCA for variable selection for high-dimensional genotype data coded 0, 1, or 2 based on the reference allele counts. These authors ranked the original variables based on their weights in the first two PCs. These approaches, however, are arguably too arbitrary to be used for general purpose variable selection.

In this study we propose to use PCA for variable selection in a binary classification problem or a case-control study. We achieve this by using data from one class to fit a PCA model. The estimated parameters thus obtained are used to reconstruct the data from another class. The variables that are not reconstructed accurately are considered to be important predictors to distinguish between the two classes. The method is effective even for small sample sizes. The results, however, could be improved by accurately estimating the parameters in the training phase. Therefore, we use the class with the larger sample size for training so that the estimated parameters are as accurate as possible. In some cases it may be easier to acquire a larger sample size for one class compared to the other. For example, healthy subjects may be abundant while the diseased subjects may be rare. Training of a classification technique, such as penalized logistic regression and random forest, usually requires a sufficiently large number of samples in each class. In our case, in such case-control association studies we could use the data from healthy subjects for fitting a PCA model. A simple procedure to obtain p-values corresponding to each variable is also proposed since such a measure is relatively easier to interpret for decision making purposes.

For illustration we focus on a genomics problem, emphasizing that our method is much more widely applicable. We evaluate our method through an extensive simulation study and consider its performance in a semi-real gene expression study where the true differentially expressed genes are known. We further evaluate the method through application to two publicly available high-dimensional datasets: a childhood acute lymphoblastic leukemia gene expression dataset and a genome-wide association study dataset. Our method is straightforward to understand, computationally efficient, easy to implement and empirical evaluation on synthetic and several real-world datasets suggests that it is more effective in variable selection compared to existing methods.

## Methods

PCA is traditionally used as a dimension reduction technique and is a useful tool to visualize high-dimensional data in a manageable low-dimensional space [19, 20, 21, 22, 23, 24, 25]. It uses an orthogonal transformation to transform a set of *p* (possibly correlated) observed variables into another set of *p* uncorrelated variables termed principal components (PCs). PCs are uncorrelated linear functions of the originally observed variables that successively maximize variance such that the first PC stands for the axis along which the observed data exhibit the largest variance; the second PC stands for the axis that is orthogonal to the first PC and along which the observed data exhibit the second largest variance; the third PC stands for the axis that is orthogonal to the first two PCs and along which the observed data exhibit the third largest variance, and so on. In this way, the *p* orthogonal dimensions of variability that exist in the data are captured in *p* PCs and the proportion of variability that each PC accounts for accumulates to the total variation in the data. The sole objective of PCA is to capture as much variation as possible in the first few PCs. It is, therefore, often the case that the first *q* (*q* ≪ *p*) PCs retain conceivably useful information in the observed data and the rest contain variation mainly due to noise [22].

More formally, let *y_ij_* denote a real-valued observation of the *j*th variable made on the *i*th subject, where *i* = 1,…, *n* and *j* = 1,…, *p*. Assume that the *n* observations are arranged in *n* rows of an *n × p* data matrix *Y* with columns corresponding to *p* variables or features. We standardize columns of *Y* to have zero mean and unit standard deviation, and store the resultant values in a data matrix *X*, that is, the elements *x_ij_* of *X* are obtained by 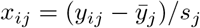, where 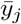 and *s_j_* are the mean and standard deviation of the *j*th column of *Y*, respectively. The PCA can be performed by singular value decomposition (SVD) of *X*; that is, the *n* × *p* matrix *X* of rank *r* ≤ min(*n, p*) is decomposed as

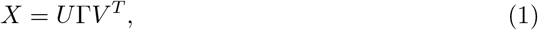

where *U* is a *n* × *r* orthonormal matrix (*U^T^U* = *I_r_*), Γ is a *r* × *r* diagonal matrix containing *r* non-negative singular values in decreasing order of magnitude on the diagonal and *V* is a *p* × *r* matrix with orthonormal columns (*V^T^V* = *I_r_*). Denote the sample correlation matrix of *X* by *R*, which can be expressed as

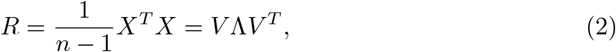

where Λ = 1/(*n* − 1)Γ^2^ is a *r* × *r* diagonal matrix containing *r* non-zero positive eigenvalues ***λ*** = (*λ*_1_,…, *λ_r_*)^*T*^ of matrix *R* on the diagonal in decreasing order of magnitude. It follows that the *r* columns of matrix *V* contain the eigenvectors of *X^T^X* and hence are the desired directions of variation. The derived set of *r* transformed variables (the PCs) are obtained by

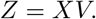

It is important to note that the matrix *U* above contains standardized PCs in its columns and is a scaled version of *Z*, which is provided additionally in (1). To see this, multiply (1) on the right by *V* to obtain

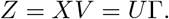

In practice, the first *q* ≪ *r* major components are of greater interest since they account for most of the variation in the data. Without undue loss of information, the dimension of *Z* may thus be reduced from *p* to *q*, that is

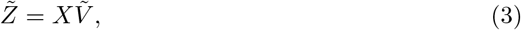

where 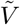 is a *p* × *q* matrix that contains the first *q* columns of *V* and 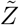 contains the first *q* PCs in its columns. The set of first *q* PCs is a lower dimensional representation of a p-dimensional dataset and can be used to uncover trends and patterns in the data, which is probably the most prevalent application of PCA. In addition, the most informative PCs can be used in subsequent analyses of the data [for example, see 26]. However, there are concerns with this approach. First, like the *p* original variables, the *q* PCs (although informative because they account for most of the variation in the data) are not easy to interpret [15, 27] and often misleading or imaginary interpretations are associated with PCs [22]. Second, in high-dimensional datasets the level of noise due to high-dimensionality often obscures useful reproducible patterns (true structure rather than random association) in the data [28]. Third, the PCs are developed in an unsupervised manner so may not be optimal for further analyses such as regression. In such cases it is assumed that only a small number of variables are able to uncover interpretable patterns in the observed data and variable selection (rather than extraction) methods are usually preferred.

A lower rank approximation of *X* can be obtained by

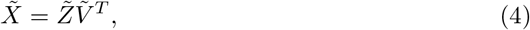

which is the best approximation of *X* in the least-squares sense by a matrix of rank *q* [29, 30]. The value of *q* is chosen using available heuristic options such as a plot of eigenvalues (or singular values) in decreasing order of magnitude, known as a scree-plot or the cumulative proportion of explained variance (CPEV), which chooses a smallest value of *q* for which the CPEV exceeds a pre-specified quantity *φ* that takes a value between 0 and 1. Commonly used values of *φ* range from 0.7 to 0.9 [22]. A value of *φ* = 0.8 means that the chosen number of components *q* explains at least 80% of the total variance in the observed data.

PCA is computationally efficient and scalable to high-dimensional datasets. In this paper, we adapt PCA to variable selection in a classification problem—rather than its traditional use of variable extraction and thereby visualization. We achieve this by considering the error induced by the approximate reconstruction of an actual observation *x_ij_* using the expression in (4); that is, the error is defined by

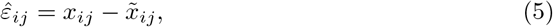

where 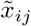 are the values contained in the *i*th row and *j*th column of 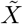. We store the elements 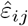 in an *n* × *p* residual matrix 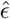.

In a case-control study, assume that *Y^−^* is an *n*_1_ × *p* data matrix containing the data from the *n*_1_ control samples and *Y* ^+^ is an *n*_2_ × *p* data matrix containing observations from the *n*_2_ cases. We standardize the *p* columns of *Y^−^* to have zero mean and unit standard deviation, and store the resultant values in a matrix *X^−^*, that is, the elements 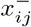 of *X^−^* are obtained by 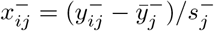, where 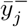 and 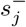 are the mean and standard deviation of the *j*th column of matrix *Y^−^*, respectively. We decompose *X^−^* as given in (1) and estimate the parameter *V^−^*. We also standardize the columns of *Y* ^+^ using the means and standard deviation of the corresponding columns in *Y^−^* and store the resultant values in *X*^+^; that is, the elements 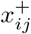 of *X*^+^ 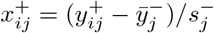. We reconstruct *X*^+^ by substituting the estimated parameter *V^−^* in (4). The reconstruction error of *X*^+^ is computed using the expression given in (5) as

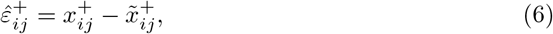

where 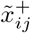 is the low rank approximation of 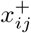. If a variable is differentially expressed between *X^−^* and *X*^+^, this will be reflected in the reconstruction error. The larger the reconstruction error the larger will be the discriminative ability of that particular variable. We compute the mean of the *j*th column of 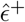 and denote it by 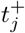; that is,

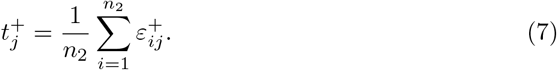

If a variable in *X*^+^ is generated by a different model then its corresponding 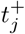 will follow a different distribution. In high-dimensional problems only a small subset of variables usually turns out to be useful in terms of their association with a response variable of interest. In such situations the realizations of 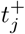 associated with the majority of unimportant variables will behave similarly in distribution. The differential expression of variables in two groups or the class prediction ability of a variable will be reflected by its associated value of 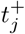. In other words, the values of 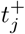 that correspond to variables that are differentially expressed between the two groups will deviate more from the center—zero—of the distribution of 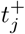. The further away a 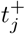 is from zero, the better its associated variable would perform as a predictor in classification. One can thus rank the *p* variables based on the magnitude of the statistic 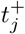 from most important to least important.

One can even calculate p-values associated with 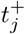 if the elements of 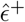 are assumed to be independent and distributed according to the Gaussian distribution; that is,

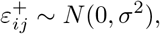

where *σ*^2^ is a variance that needs to be estimated. This is a common assumption generally made when inference procedures are developed for a PCA model [for example, see 31]. If *X^−^* and *X*^+^ are generated by the same model then the distribution of 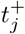 is easy to characterise; that is,

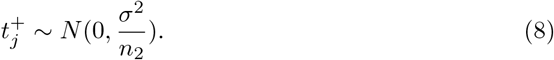

This turns a variable selection problem to an outlier identification problem and a variable that is associated with an outlying 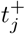 will have discriminative ability to differentiate between groups when used in a predictive model such as logistic regression.

Note that using the sample variance as an estimator of *σ*^2^ may obscure the outlying values of 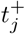 since the sample variance is not robust against outliers. It is important to use a robust estimate of *σ*^2^ such as mean absolute deviation (MAD), which has a 50% breakdown point [32] as opposed to sample standard deviation, which has a 0% breakdown point. As noticed above, in high-dimensional problems one usually does not expect to have a large number of significant variables (since MAD breaks down if the proportion of significant variables is larger than 50%). The MAD estimator of *σ* is

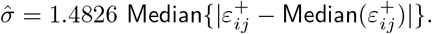

The above procedure could be used to identify potential discriminatory variables even if we have a single case sample (many healthy baseline observations vs. a single case). It could be particularly useful when one wishes to identify differentially expressed genes in a gene expression profile of a small number of available patients. In some situations it may be that some of the samples in the case group are different with respect to some variables even though they all belong to a single group. For example, one particular type of leukaemia might have some unknown subtypes and identifying subtypes of a disease could be useful in precision medicine and personalized treatments [33, 34]. The one-by-one analysis makes it possible to further identify those variables which are differentially expressed and are common (or uncommon) to all patients in the same group. This makes our procedure different and more advantageous from the variable selection procedures that are commonly used in practice.

In the rest of this document we refer to our PCA based approach as Variable Selection based on PCA (VSPCA) and formalize it in the following algorithm:

1. Obtain *X^−^* by standardizing the data matrix *Y ^−^*; that is, 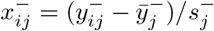.
2. Estimate the parameter *V ^−^* using the control sample *X^−^* and the expression in (1).
3. Obtain *X*^+^ using 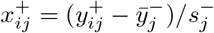.
4. Use the proportion of total variance explained criterion or scree-plot to choose *q* and calculate 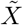 using the estimates 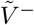 and the expression in (4).
5. Compute 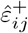 using (6).
6. Estimate *σ* using a robust MAD estimator given by

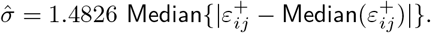
7. Calculate 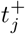 using (7).
8. Calculate p-values under the assumption that 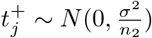. A smaller p-value indicates the inconsistency of associated variable in *X*^+^ with the one in *X^−^* and hence is the important variable.

Note that in some circumstances one may wish to reduce the number of variables from *p* to a smaller number *k* (i.e., to filter out the *p* − *k* least important variables). This is also useful when the number of variables is small, in which case one does not have enough data to estimate *σ*, or when the required normality assumption for the error is not valid. In such cases step-8 of the algorithm can be dropped and variables can be ranked based on the relative magnitude of 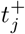. Those variables that correspond to the *k* largest magnitudes of 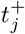 can be selected as the superset of predictors. Note also that the p-values obtained via VSPCA need to be adjusted for multiple testing. Unless otherwise specified, all p-values reported in this paper are adjusted for multiple testing via the ‘fdr’ method [35] implemented in the p.adjust() function of the R ‘stats’ package.

We recommend examining the quantile-quantile plot (qq-plot) of 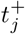 and the histograms of unadjusted p-values for diagnostic purposes. The distribution of 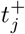 is expected to be normally distributed and in a typical high-dimensional problem where a large number of variables are irrelevant, the distribution of 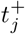 should be normal at least at the center. The heavier tails of the distribution in the qq-plot indicate significant variables. Similarly, under the null hypothesis of no relevant variable, one would expect the two-tailed p-values to be uniformly distributed between 0 and 1. If there are informative variables, this would be indicated in the histogram by a sharp spike near 0. The rest of the shape is still expected to be uniform.

## Simulation study

We used simulations to study the empirical performance of our method and to explore the circumstances in which it substantially dominates some state-of-the-art methods. We considered a simulation setting similar to the one created in [36] and later adopted in [37]. A range of simulation scenarios was considered based on combinations of the following four factors: (i) size of the training dataset *Y ^−^*, *n*_1_ ∈ {10, 20, 30, 50, 100, 200}; (ii) size of the test dataset *Y*^+^, *n*_2_ ∈ {10, 20, 30}; (iii) proportion of explained variance in the training data, *φ* ∈ {0.2, 0.4, 0.6, 0.8} that in turn determines the number of components (*q*) to be used for reconstruction of the test dataset; (iv) the strength of correlation between variables, *ρ* ∈ {0, 0.5, 0.8}. A sample of size *n*_1_ was generated from a *p* = 3000 dimensional normal distribution with mean vector *μ*_1_ = 0_*p*_ and a covariance matrix 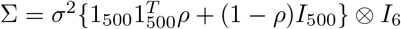 where 0_*p*_ is a column vector of 0’s with dimension *p* × 1, 1_500_ is a column vector of 1’s with dimension 500 × 1, *I_d_* is an identity matrix with dimension *d* × *d*, ꕕ denotes the Kronecker product and *σ*^2^ = 1. The test sample of size *n*_2_ was obtained in a similar way as the training dataset except that the means of 600 randomly chosen variables were changed by *δ* ∈ {±0.6, ±0.9, ±1.2}, where each of the six values of *δ* had the same chance of selection. For each combination of the four factors, we conducted 1000 independent simulation experiments.

One of the most powerful univariate statistical tests is the empirical Bayes moderated t-test [1], which is a popular choice for the analysis of data from microarray experiments [38]. The test borrows information between genes through shrinking gene-specific variances towards the global variance and is implemented in the ‘limma’ package of R [39] (a reason for it is sometimes referred to as LIMMA.) We chose this test for comparison purposes and will refer to it as MTT for the rest of the paper.

The full set of simulation results is provided in supplementary materials (see Fig S1-S18). The results for *n*_1_ ∈ {10, 30, 100}, *n*_2_ = 10, *ρ* = 0.8, *φ* ∈ {0.2, 0.4, 0.6, 0.8} are given in Fig 1 and 2. Fig 1 is focused on empirical power and FDR evaluated at a fixed nominal level of significance *α* = 0.01. The VSPCA maintains lower FDR than MTT in all simulation scenarios shown in Fig 1 (see the results for *δ* ≠ 0) and is much powerful than MTT (see the results for *δ* ≠ 0 in Fig 1). Fig 2 shows top-ranked variables in terms of their predictive importance and the proportions of true nulls among them. In general, for moderate (see Fig S7-S12) to highly (see Fig 2 and Fig S13-S18) correlated variables the VSPCA performed substantially better than MTT in terms of both power and FDR. For uncorrelated (see Fig S1-S6) to weakly (results not shown) correlated variables, the MTT performed slightly better when *n*_1_ was small; otherwise, both methods performed almost equally well (VSPCA with a suitable value of *φ*).

**Fig 1.**
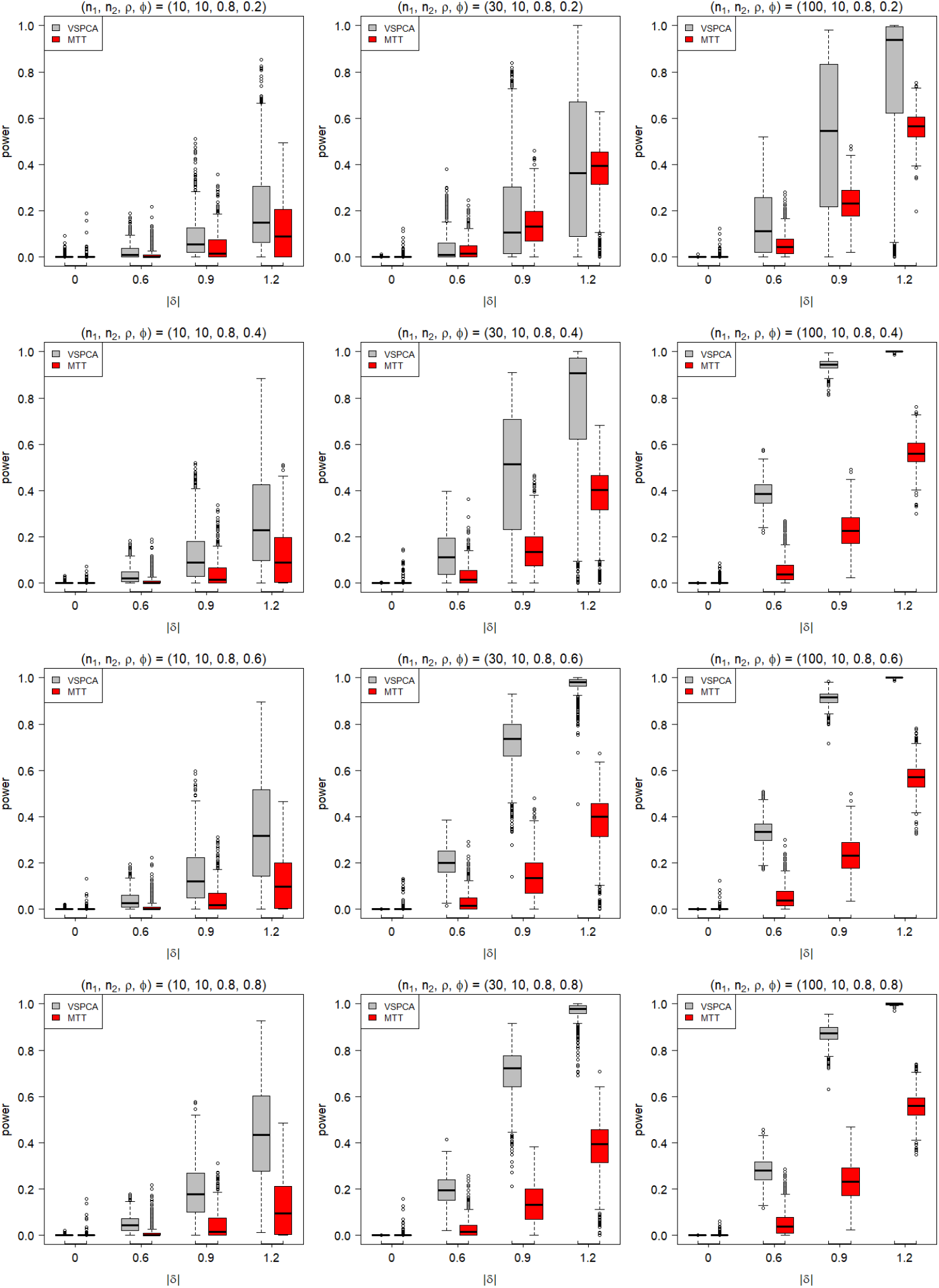
Empirical power of VSPCA and MTT to correctly detect differences between groups (differentiate zero and non-zero values of *δ*). Plotted are the proportion of adjusted p-values that were less than the nominal level of significance *α* = 0.01 for absolute values of *δ*. We set *ρ* = 0.8 and *φ* ∈ {0.2, 0.4, 0.6, 0.8}. Results in each graph are based on 1000 simulated datasets.

**Fig 2.**
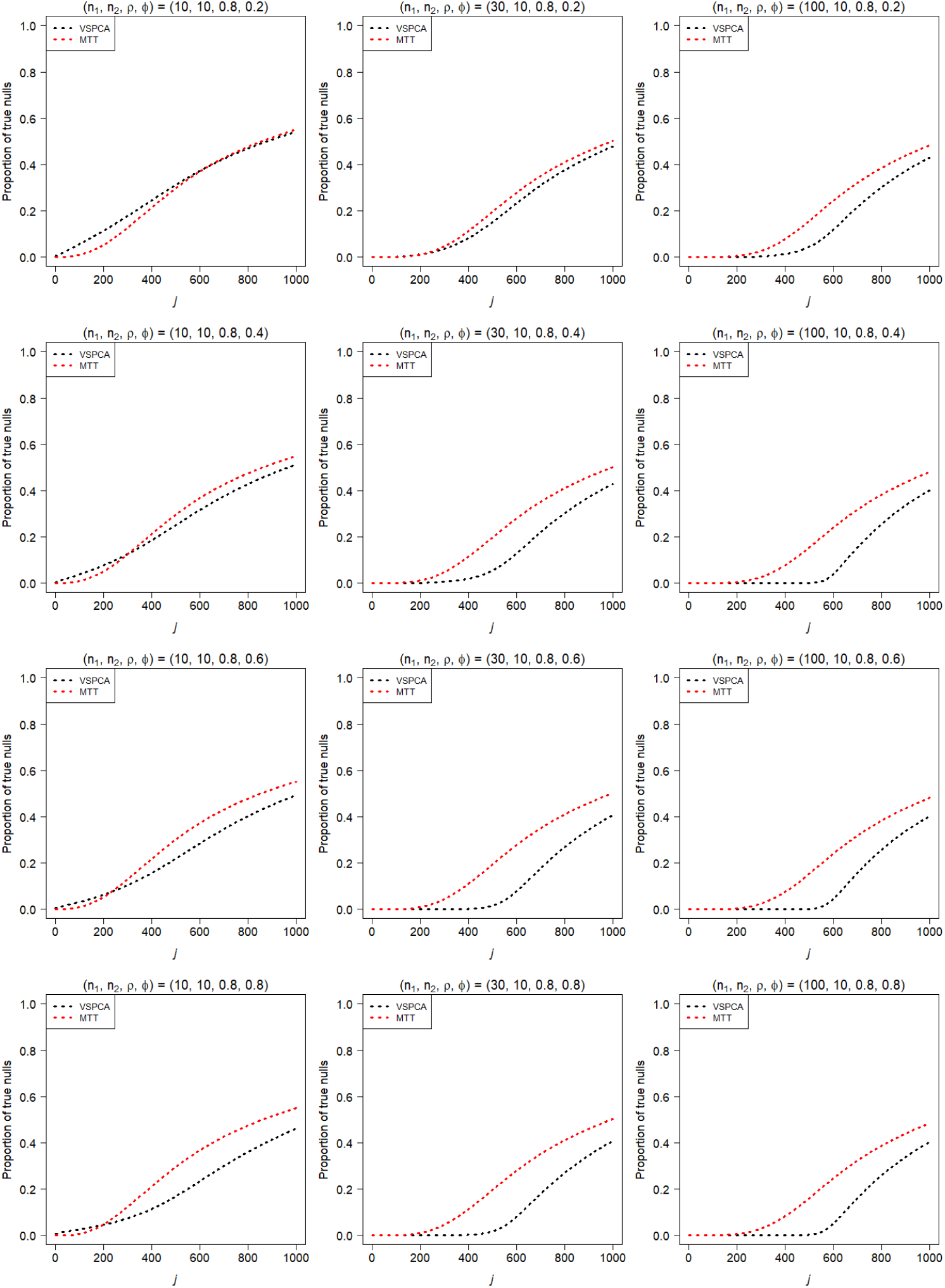
Shown in each plot are the proportions of true nulls among the *j* top-ranked variables with the ability to differentiate between groups (*j* = 1, 2,…, 1000). Results in each graph are averaged over 1000 simulated datasets. The simulation setup is as for Fig 1.

Here we summarize the key characteristics of VSPCA we found through our experience with a variety of simulation experiments including those provided here and in supporting information:

*n*_1_: The power of VSPCA vastly improved with increase in *n*_1_.
*n*_2_: The increase in *n*_2_ had a very small impact on the performance of VSPCA.
*ρ*: The correlation between variables in the dataset played a vital role. The stronger the correlation between variables in the dataset, the higher the power of VSPCA and lower the FDR because the method gains power and reduces FDR through taking into account the correlation between variables.
*φ*: A suitable value of the factor *φ* depended on the values of *n*_1_ and *ρ*. A smaller value of *φ* was required if a larger sample was used to train the PCA model under a moderate to strong value of *ρ* (see Fig 1). For smaller values of *n*_1_ and *ρ* a larger value *φ* produced good results.

## Applications

### Analysis of Platinum Spike Dataset

We test and illustrate our new variable selection procedure by applying it to the Platinum Spike [40] dataset. This is one of the largest semi-real datasets for which the true differentially expressed genes are known and is accessible through GEO Series accession number GSE21344. Hence, it has been widely used for benchmarking microarray analysis methods [for example, see 41, 42, and references therein].

The dataset is produced by a controlled experiment [40] with two experimental conditions (condition-A and condition-B) and nine (three biological × three technical) replicates per condition. Gene expression data were obtained with Affymetrix Drosophila Genome 2.0 microarrays. The dataset has a total of 18,769 probe sets (excluding Affymetrix internal control probes), of which 5,587 were spiked-in. Among these, 1,940 (34.7%) were differentially expressed (known positives) to varying degrees between 1.2- and 4-fold (1,057 over-expressed, 883 under-expressed), while the remaining 3,406 (61.0%) were spiked-in at the same concentration in both conditions (known negatives).

Although the sample size of the training set (whichever of condition-A and condition-B one wishes to choose as a training set of data) is very small (*n*_1_ = 9), this dataset is worth considering here because of its benchmark status as described above [43]. The analysis of this dataset will help understand the performance of VSPCA under this common scenario of a very small sample size.

We normalized the dataset using the Affymetrix MAS5 algorithm implemented in the *mas5* function of the R package ‘affy’ [44]. We applied our variable selection procedure to the processed data. The PCA model was trained on data produced under condition-B. The scree-plot is shown in Fig 3a and the CPEV plots is given in Fig 3b. The scree-plot showed a mild elbow at *q* = 3. These three components together explained 42.9% of the total variation in the training data. We set *φ* = .7 to exclude three components for the purpose of denoising (the variance explained by component-9 is zero), which led us to choose *q* = 6 to obtain the low-rank approximation of the data produced under condition-A. For comparison purposes we also presented the results obtained from the MTT.

**Fig 3.**
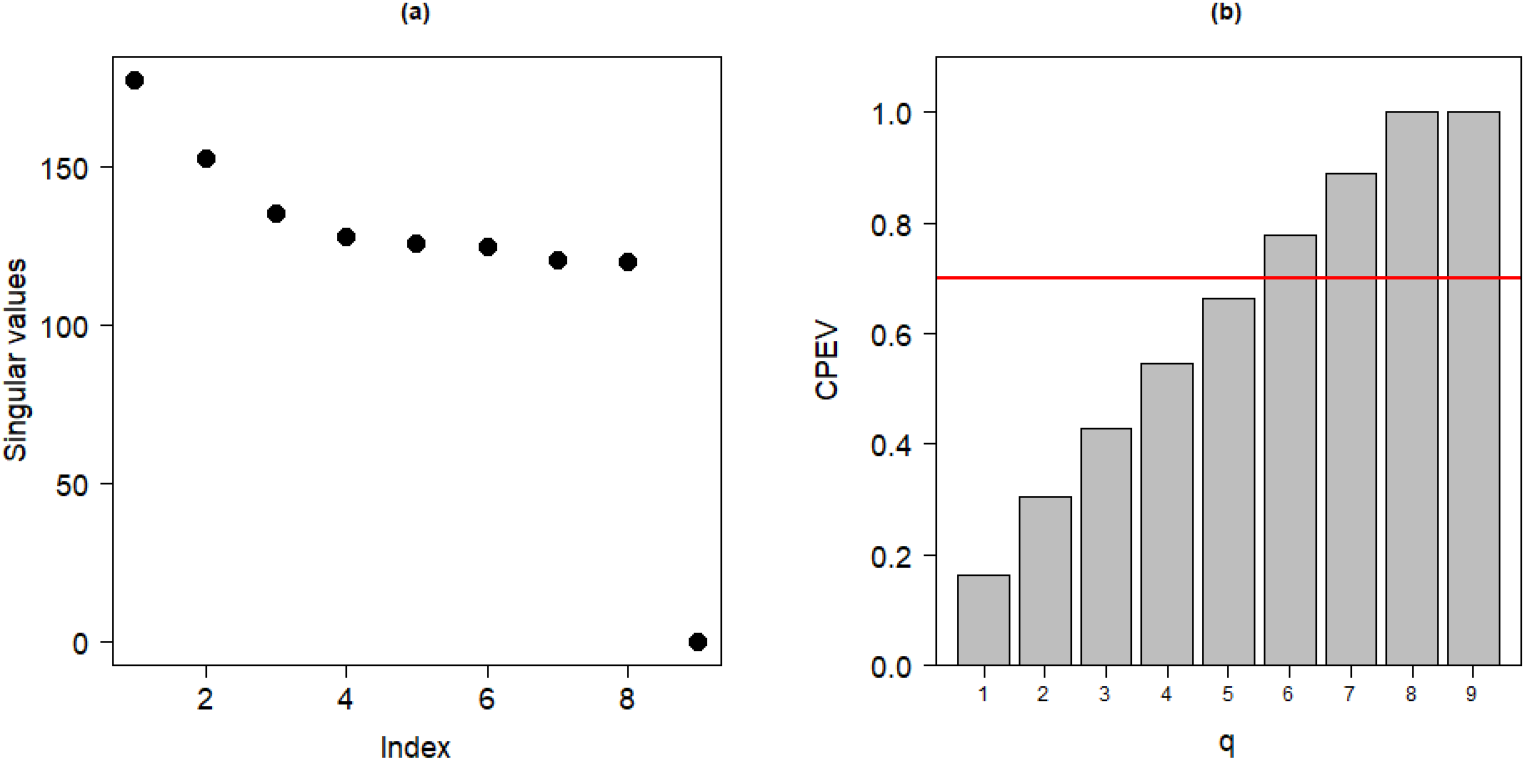
The plot in (a) shows the scree plot for the training data (9 samples produced under condition-B) of Platinum spike dataset and the plot in (b) shows the cumulative proportion of explained variance (CPEV) for each value of *q* with the horizontal line in red indicating *φ* = .7.

The qq-plot is given in Fig 4 with heavy tails indicating a large number of differentially expressed genes between the two experimental conditions. The VSPCA led to a lower False Positive Rate (FPR) (see Fig 5a) in comparison to MTT (see Fig 5c). The two methods, however, performed almost equally well in identifying true differentially expressed genes between the two experimental conditions (see Fig 5b and 5d). The p-values of VSPCA are shown in Fig 6 in comparison to the p-values obtained using MTT. The p-values of VSPCA for the true negatives were much larger than the p-values based on MTT (see Fig 6a) while the p-values of VSPCA for true positives were much smaller compared to the p-values of MTT (see Fig 6b). In addition, for the true positives the VSPCA further demarcated the genes that have the highest discriminative ability (the super predictors). Based on the ROC curve in Fig 7a and Fig 7b in which we plot the proportion of true nulls among the top-ranked differentially expressed genes calculated at various cut-off points, our multivariate approach appeared to have performed almost as well as the univariate MTT.

**Fig 4.**
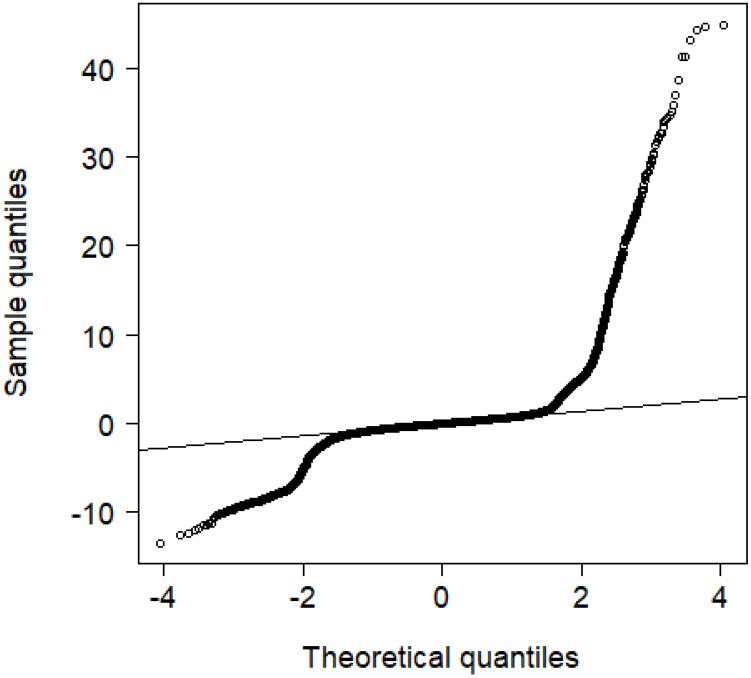
Platinum spike dataset. Quantiles of 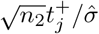 against the quantiles of standard normal distribution.

**Fig 5.**
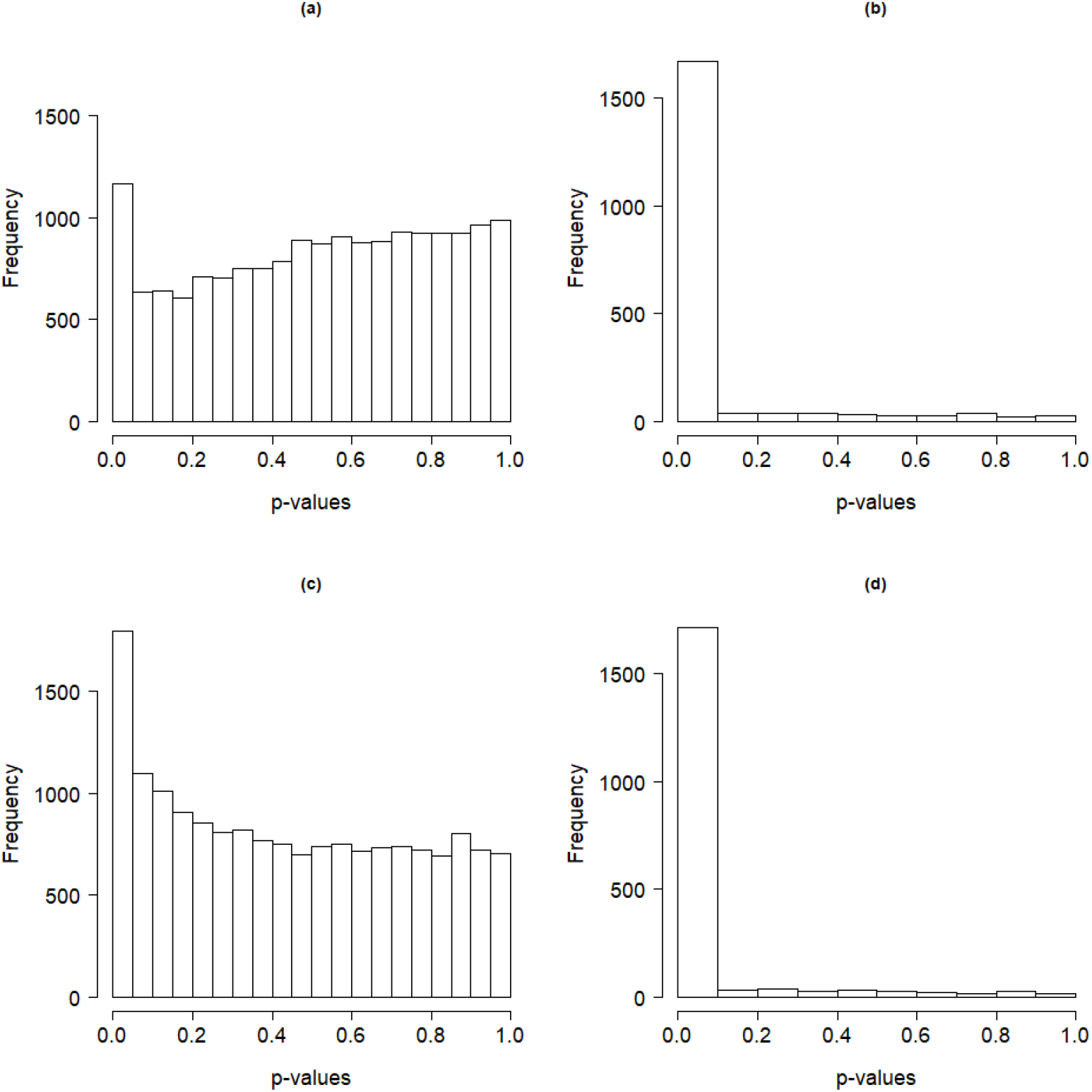
Histograms of unadjusted p-values for Platinum spike dataset: p-values calculated for (a) known negatives using VSPCA, (b) known positives using VSPCA, (c) known negatives using MTT, and (d) known positives using MTT.

**Fig 6.**
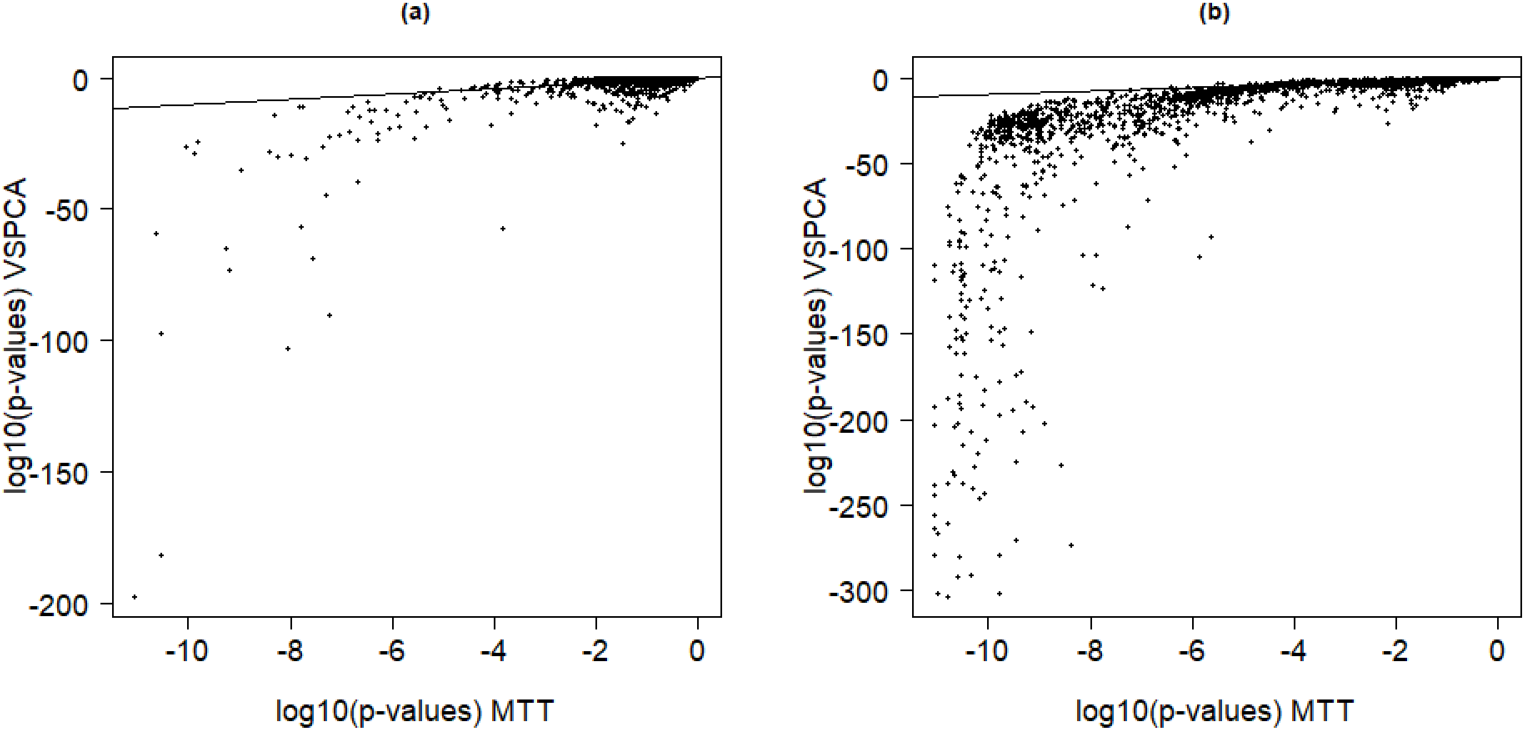
Logarithm of p-values obtained using VSPCA against logarithm of p-values obtained using MTT for Platinum spike dataset: (a) p-values for known negatives (b) p-values for known positives. The equality, *x* = *y* values, reference lines are added.

**Fig 7.**
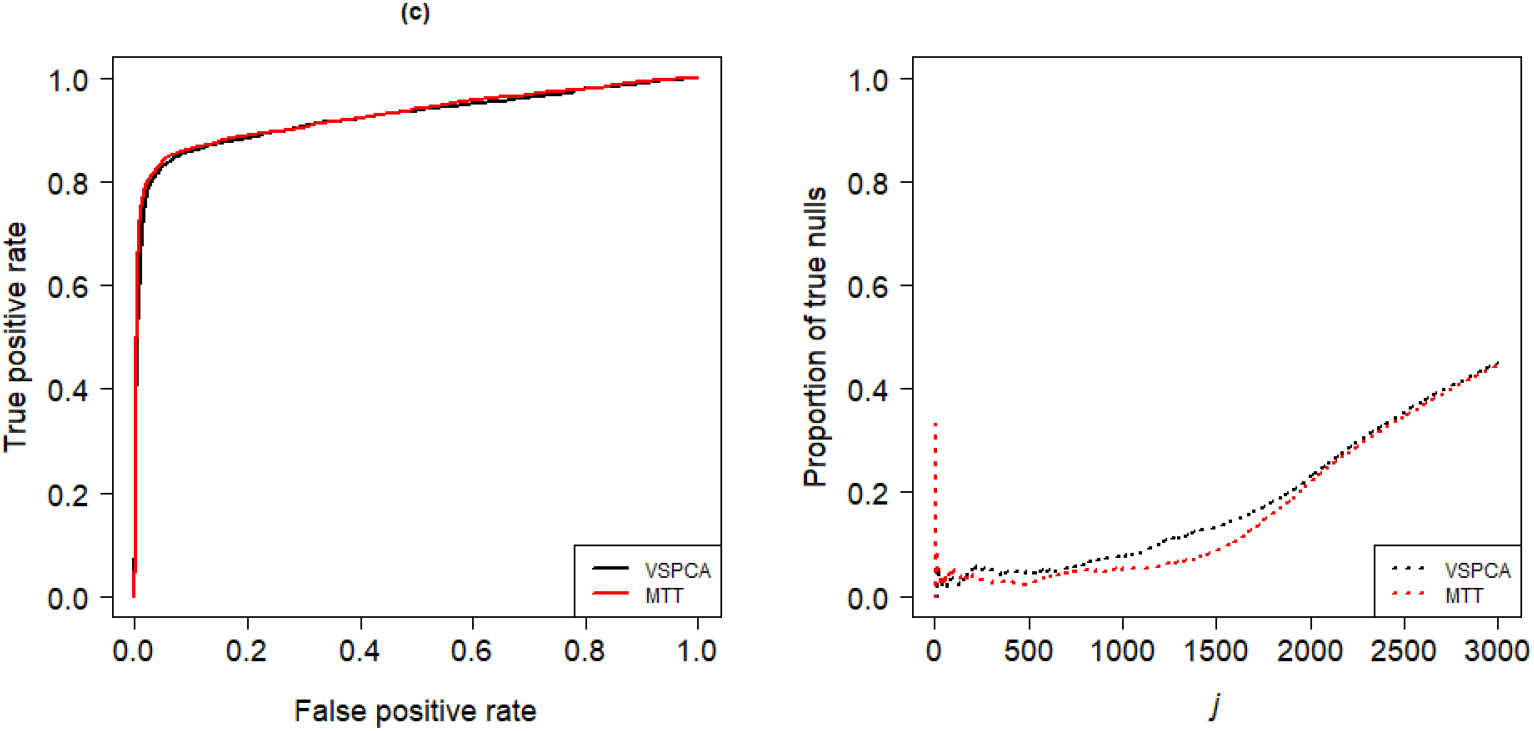
Comparison of VSPCA and MTT for Platinum spike dataset (a) ROC curve (b) False positive detected among the *j* top-ranked deferentially expressed genes (*j* = 1, 2,…, 3000).

### Childhood acute lymphoblastic leukemia (CALL) gene expression study

To provide additional evidence of the performance of VSPCA, we chose a childhood acute lymphoblastic leukemia (CALL) high-throughput gene-expression dataset that is studied in detail by [45] and accessible through GEO Series accession number GSE13425. The data had 22,283 genes and 190 samples in total. The 190 samples are from different subtypes of CALL: BCR-ABL (4 samples), BCR.ABL.hyperdiploidy (1 sample), E2A-rearranged.E (1 sample), E2A-rearranged.Esub, (4 samples), E2A-rearranged (8 samples), hyperdiploid (44 samples), MLL (4 samples), pre.B.ALL (44 samples), T.ALL (36 samples), TEL.AML1 (43 samples), and TEL.AML1.hyperdiploidy (1 sample). This allows us to evaluate the performance of the method under a variety of sample sizes.

We normalized the data using the Affymetrix MAS5 algorithm implemented in the R package ‘limma’ [39]. The normalized data were then used to rank and identify genes that are associated with the class labels using our VSPCA method and the MTT.

We used hyperdiploid subtype samples (*n*_1_ = 44) as a control group to train our PCA model. The T.ALL subtype samples (*n*_2_ = 36) were used as cases to identify genes that are differentially expressed from hyperdiploid subtype samples. A scree plot based on hyperdiploid subtype samples is shown in Fig 8, which suggested *q* = 7. However, we used the proportion of explained variance criterion which we believe protects against unreasonably smaller choices of *q*. We set *φ* = .8, which yielded *q* = 33 to reconstruct the test set of data (cases group).

**Fig 8.**
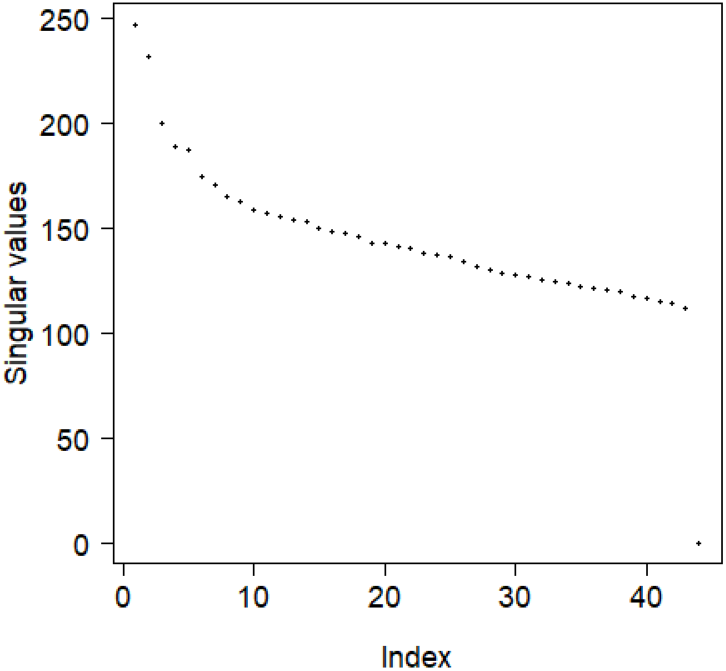
Scree plot based on hyperdiploid subtype samples of CALL dataset.

The qq-plot of 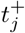 is given in Fig 9 and the histograms of unadjusted p-values are shown in Fig 10a and Fig 10b. The heavier tails of the distribution of 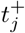 in the qq-plot indicate differentially expressed genes and the distribution seems to be normal around the center. The histograms of unadjusted p-values obtained using VSPCA and MTT revealed a large number of null genes with a spike near zero indicated a number of differentially expressed genes. The p-values obtained using VSPCA are plotted against the p-values obtained using the MTT in Fig 10c. The significant p-values obtained via VSPCA were found to be smaller in comparison to the significant values obtained via MTT. This has an advantage when it comes to correction for multiple testing; that is, the smaller a p-value, the higher the chance of remaining in the set of significant p-values once corrected for multiple testing.

**Fig 9.**
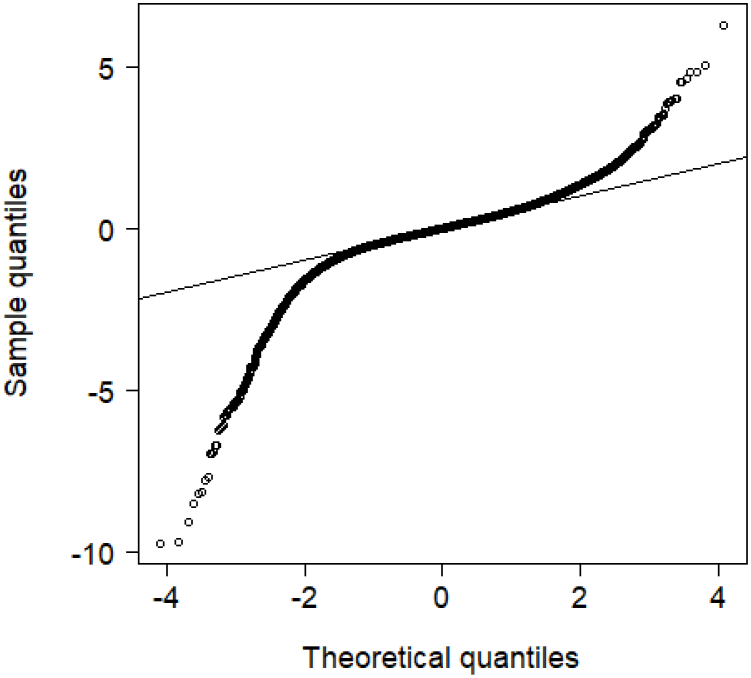
Quantiles of 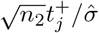 against the quantiles of standard normal distribution.

**Fig 10.**
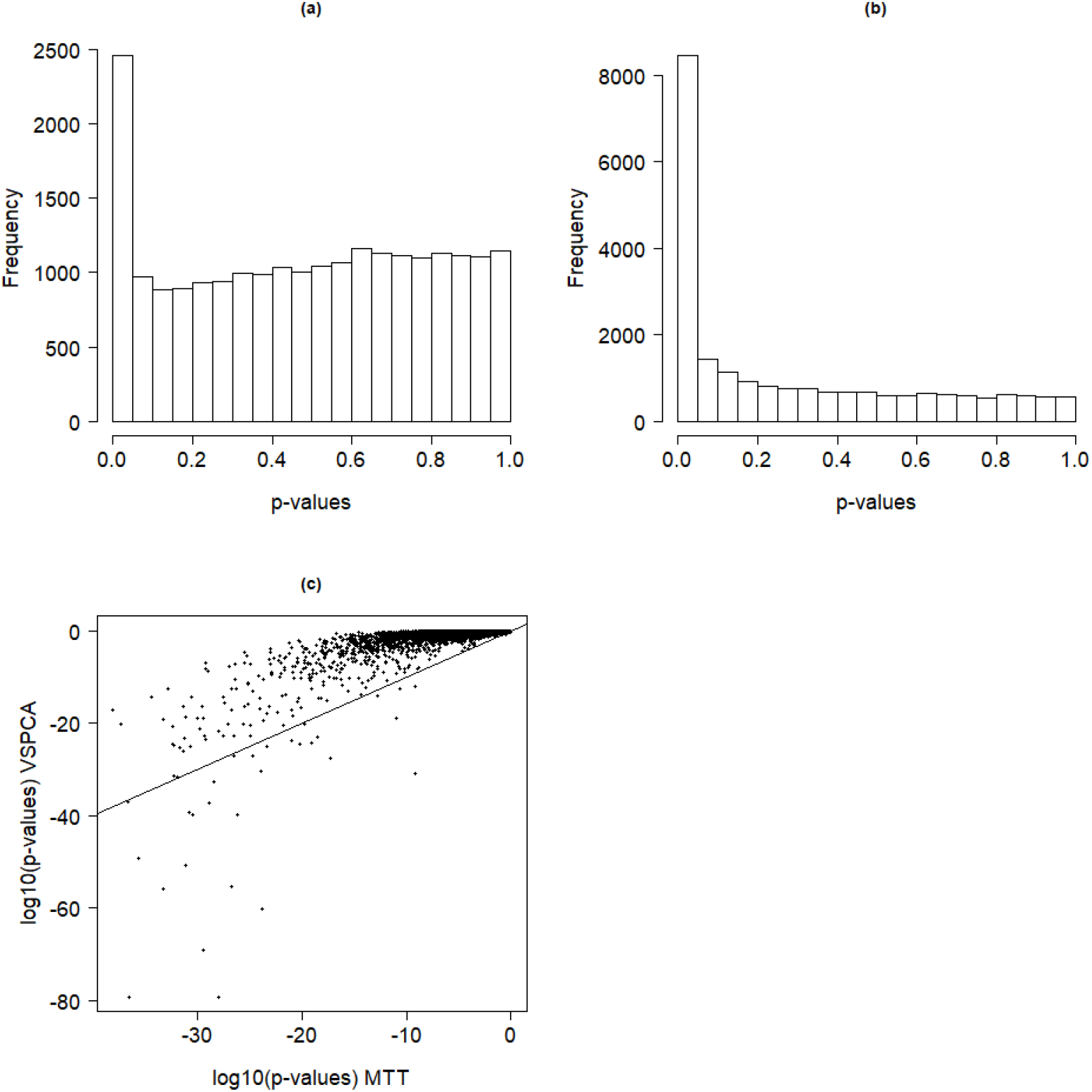
Plots of p-values: (a) Unadjusted p-values obtained via VSPCA, (b) Unadjusted p-values obtained via MTT, and (c) Logarithm of adjusted p-values obtained using VSPCA against logarithm of p-value obtained using the MTT (the equality, *x* = *y* values, reference line is added).

The MTT identified a large number of differentially expressed genes (4,980 at 0.01 level of significance) compared to VSPCA, which identified 582 differentially expressed genes. To explore the discriminative ability of the large number of additional genes identified by MTT, we performed post-selection PCA analysis using only the 582 genes selected based on VSPCA and also using the selected superset of 4,980 genes based on MTT. It turns out that the smaller superset of 582 genes selected using VSPCA has higher discriminative ability than the much larger superset of 4,980 genes selected using MTT (see Fig 11). The PCA plot based on the superset selected using VSPCA is dominated by between-group variation (see Fig 11a). It can also be observed that the hyperdiploid samples (the training data) are tightly clustered together in comparison to the T.ALL samples (the test data). This is because we took into account 33 components and dropped the rest which in general are believed to represent noise and possible outlying effects of some of the observations. This makes the procedure robust to outliers and unusual observations in the training data. In contrast, the plot based on the superset selected using MTT is dominated by within-group variation (see Fig 11b). This indicates that the MTT performance is affected by the noise, which led to a higher FPR. As expected, our new approach can substantially reduce the FPR by accounting for correlation between variables and by dropping the minor components that are often believed to stand for noise and unusual observations in the data.

**Fig 11.**
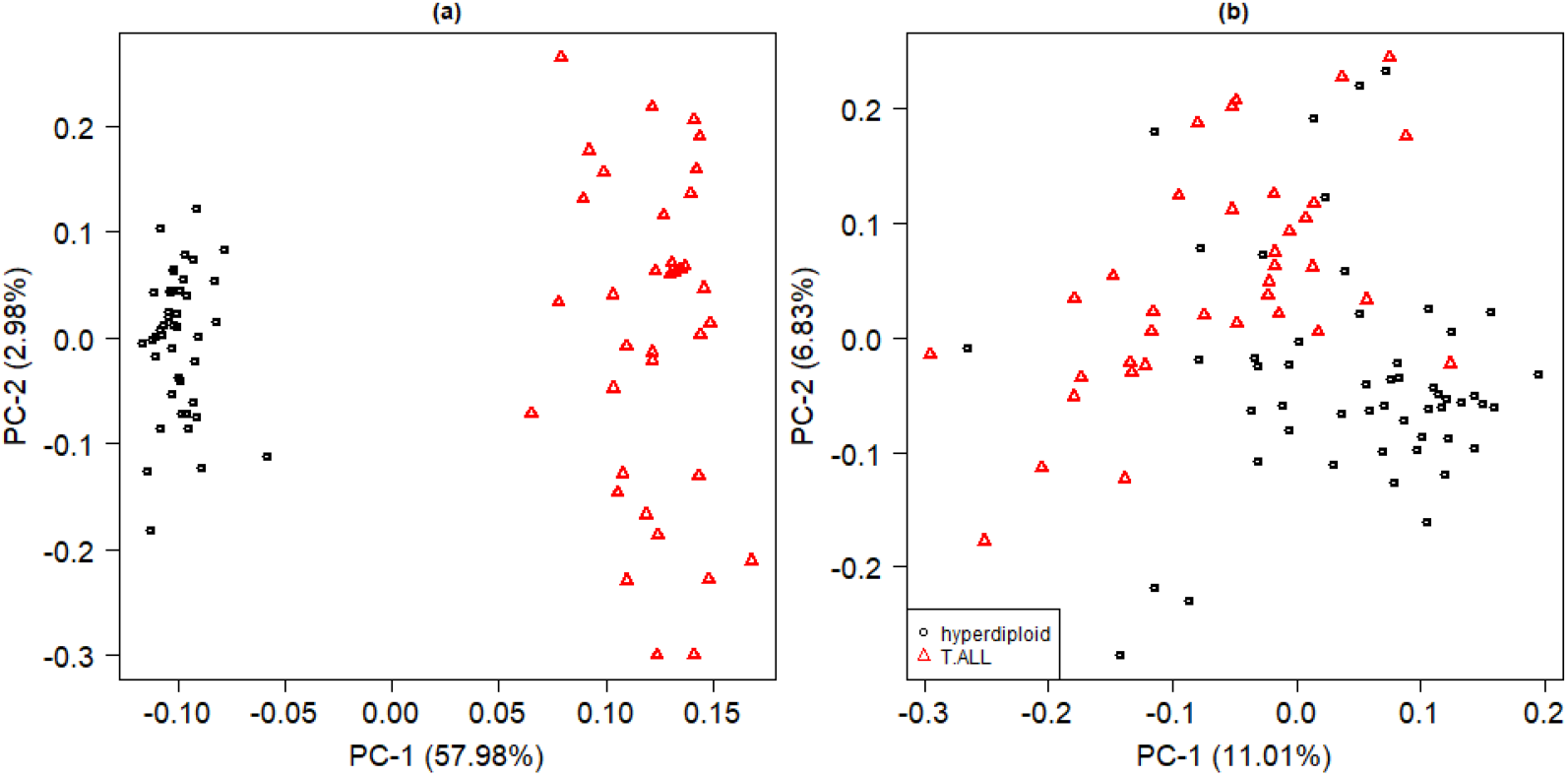
PCA plots based on the identified differentially expressed genes between hyperdiploid and T.ALL subtypes of the CALL dataset using: (a) VSPCA and (b) MTT.

To see the stability of the above results and to further test the performance of VSPCA under a smaller *n*_2_ (smaller set of test data), we re-sampled the 36 T.ALL subtype samples. As above, we used hyperdiploid samples (*n*_1_ = 44) as a training set of data and T.ALL samples as a test set of data. The value of *n*_2_ was reduced gradually from 36 to 6 by successively subtracting 5 each time. For all values of *n*_2_ < 36 that were considered here, we selected a sample without replacement from the test set of data and performed variable selection based on VSPCA and MTT using a level of significance *α* = 0.01. The experiment was repeated 1000 times for each value of *n*_2_ < 43. The variables that were significant under smaller *n*_2_ were matched with the ones that were selected under the entire available test data sample (VSPCA selected 582 genes and MTT selected 4,980 genes as above) were counted. We compared the proportions of significant variables under the entire sample size that were also significant under smaller sample sizes in more than 50% of the replicates (see Fig 12). Note that the choice of 50% (> 500/1000) is an arbitrary threshold but a similar pattern could be achieved by changing the threshold. The number of selected variables dropped much faster for MTT than it did for VSPCA as we reduced *n*_2_. With *n*_2_ = 6 the superset of 4,980 genes selected through MTT dropped by 63.5%. However, the superset of 582 genes selected through VSPCA dropped by 21.6%, which not only showed the stability of VSPCA but also showed that VSPCA could be more useful when the number of samples in the test class is smaller.

**Fig 12.**
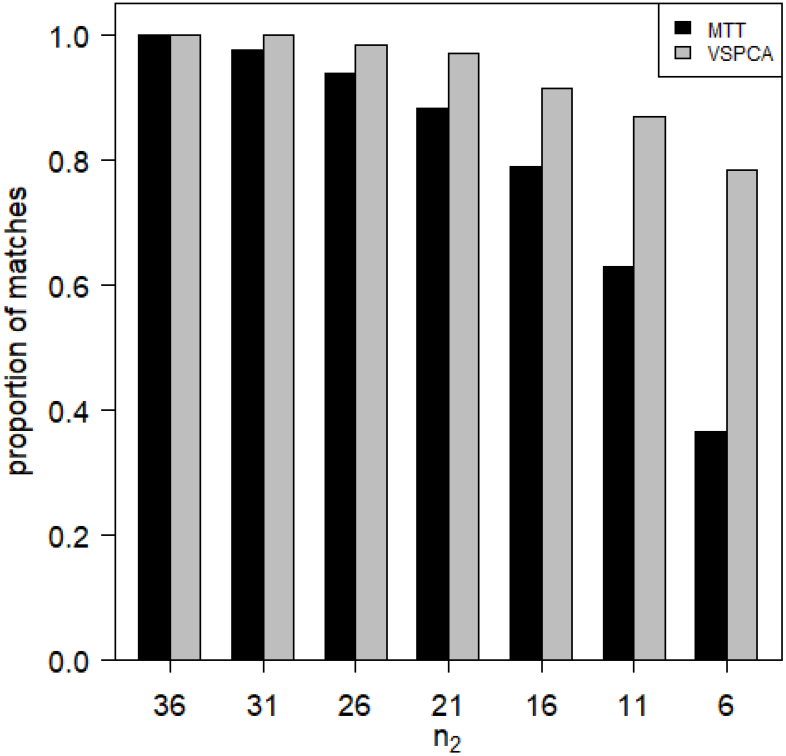
We used hyperdiploid samples (*n*_1_ = 44) as a training set of data and T.ALL samples (*n*_2_ = 36) as a test set of data. The value of *n*_2_ was reduced from 36 to 6 by successively subtracting 5 (as shown on the x-axis from left to right). For all values of *n*_2_ < 36 that are considered, we selected a sample without replacement from the test set of data and performed variable selection based on VSPCA and MTT using the nominal level of significance *α* = 0.01. The experiment was repeated 1000 times for each value of *n*_2_ < 36. The variables that turned out to be significant under smaller sample sizes and matched the ones that were selected under the entire available sample size for the test data are counted. Shown are the proportions of significant variables under the entire sample size that were also significant under smaller sample sizes in more than 50% (500/1000) of the replicates.

The VSPCA together with the post-selection PCA is useful when one is interested in identifying possible subgroups in the test data. To this end, we used the hyperdiploid subtype samples as training data and kept the training phase as it was above. However, this time we used all the rest of the subtypes as control data. Based on the VSPCA, 293 genes were found to be differentially significant at the 0.01 level of significance while the number of differentially expressed genes were 2582 when MTT was used. The PCA plots based on the identified differentially expressed genes are shown in Fig 13. The 293 genes chosen based on VSPCA reveal similarities and differences between the subtypes more clearly compared with the 2582 genes chosen based on MTT.

**Fig 13.**
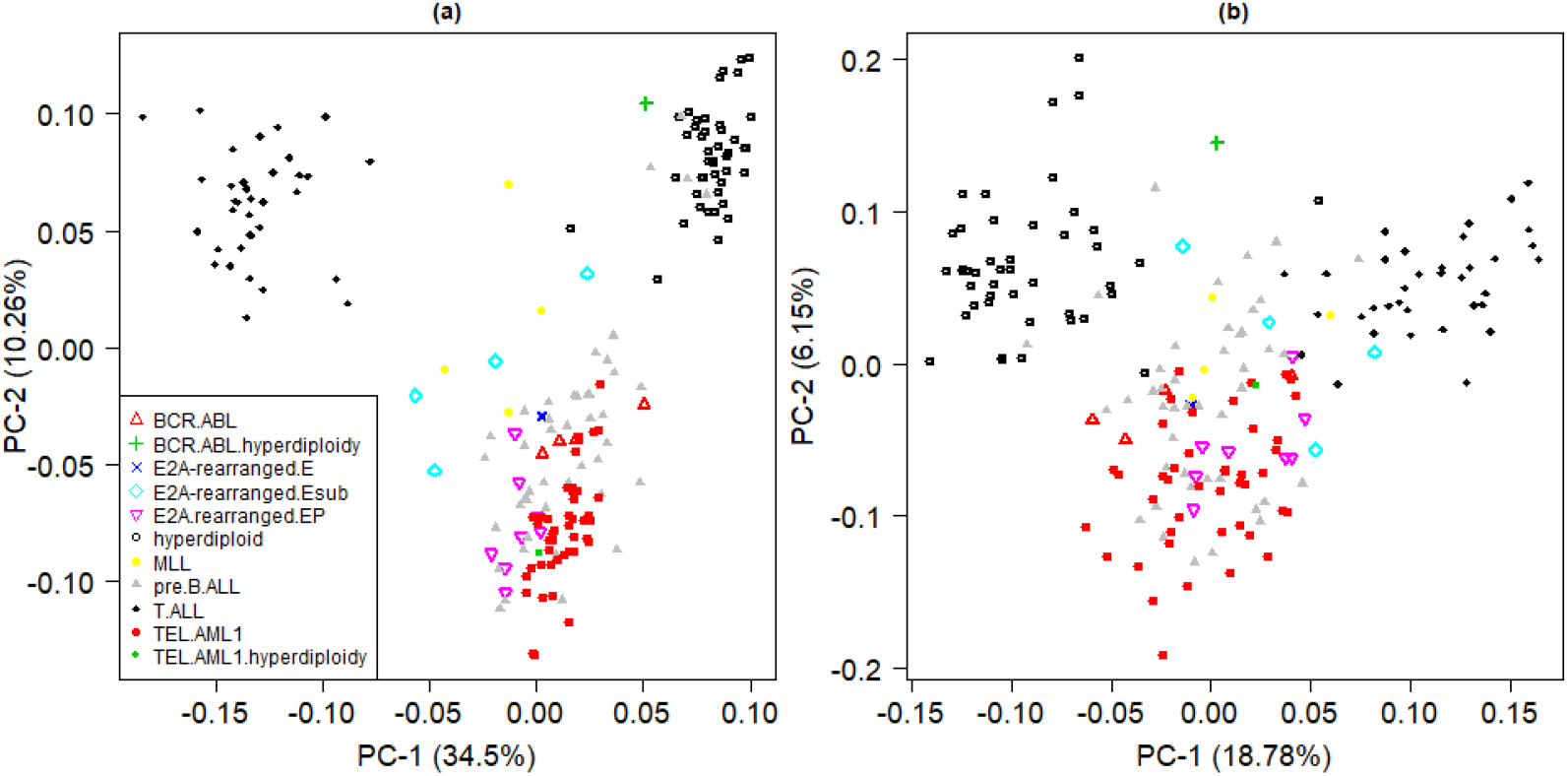
PCA plots based on the identified differentially expressed genes between hyperdiploid and the rest of the subtypes of the CALL dataset using: (a) VSPCA and (b) MTT.

To further evaluate our method, we changed the test set of data; that is, to train the PCA model we again used hyperdiploid samples but this time we used TEL.AML1 subtype (*n*_2_ = 43) as a test set of data. The number of genes that were differentially expressed between hyperdiploid subtype and TEL.AML1 subtype, at 0.01 level of significance, was 373 for VSPCA and 2196 for MTT. The MTT again identified a larger number of genes comapred to VSPCA. The histograms of unadjusted p-values are provided in Fig 14a and in Figure 14b for VSPCA and MTT, respectively. The p-values under VSPCA are plotted against the p-values under MTT in Fig 14c. As above, we performed post-selection PCA analysis using only the superset of 373 genes selected by VSPCA and also using the superset of 2,196 genes selected by MTT. The PCA analysis presented in Fig 15 shows a clear split between the observations for the two groups for both supersets. However, the VSPCA superset of 373 genes separates the two groups better (more between-group variation and less within-group variation) and therefore contains stronger predictors. The less clear separation with a much larger superset of predictors identified by MTT is evidence of higher FPR.

**Fig 14.**
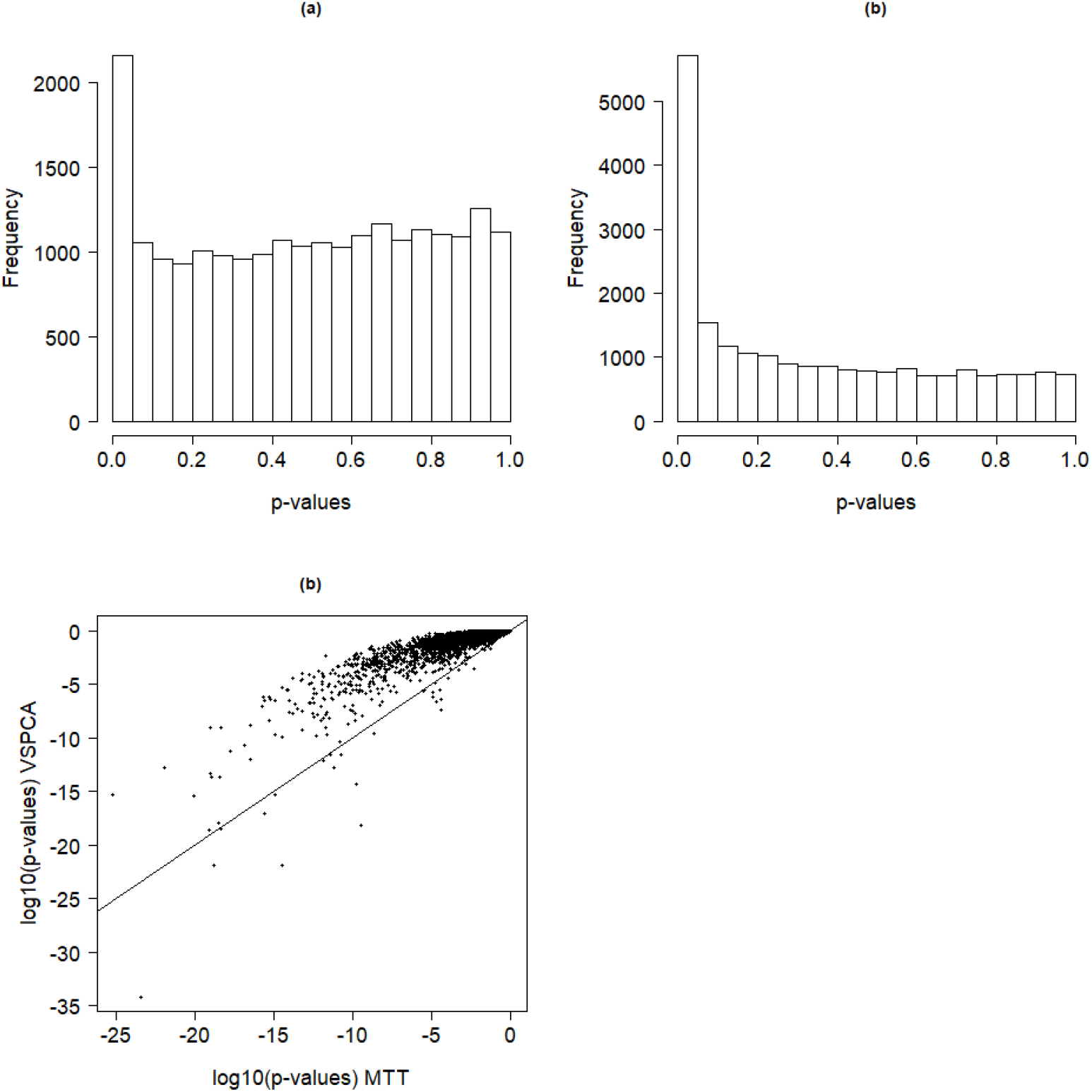
Plots of p-values: (a) Unadjusted p-values obtained via VSPCA, (b) Unadjusted p-values obtained via MTT, and (c) Logarithm of adjusted p-values obtained using VSPCA against logarithm of p-value obtained using the MTT (the equality, *x* = *y* values, reference line is added).

**Fig 15.**
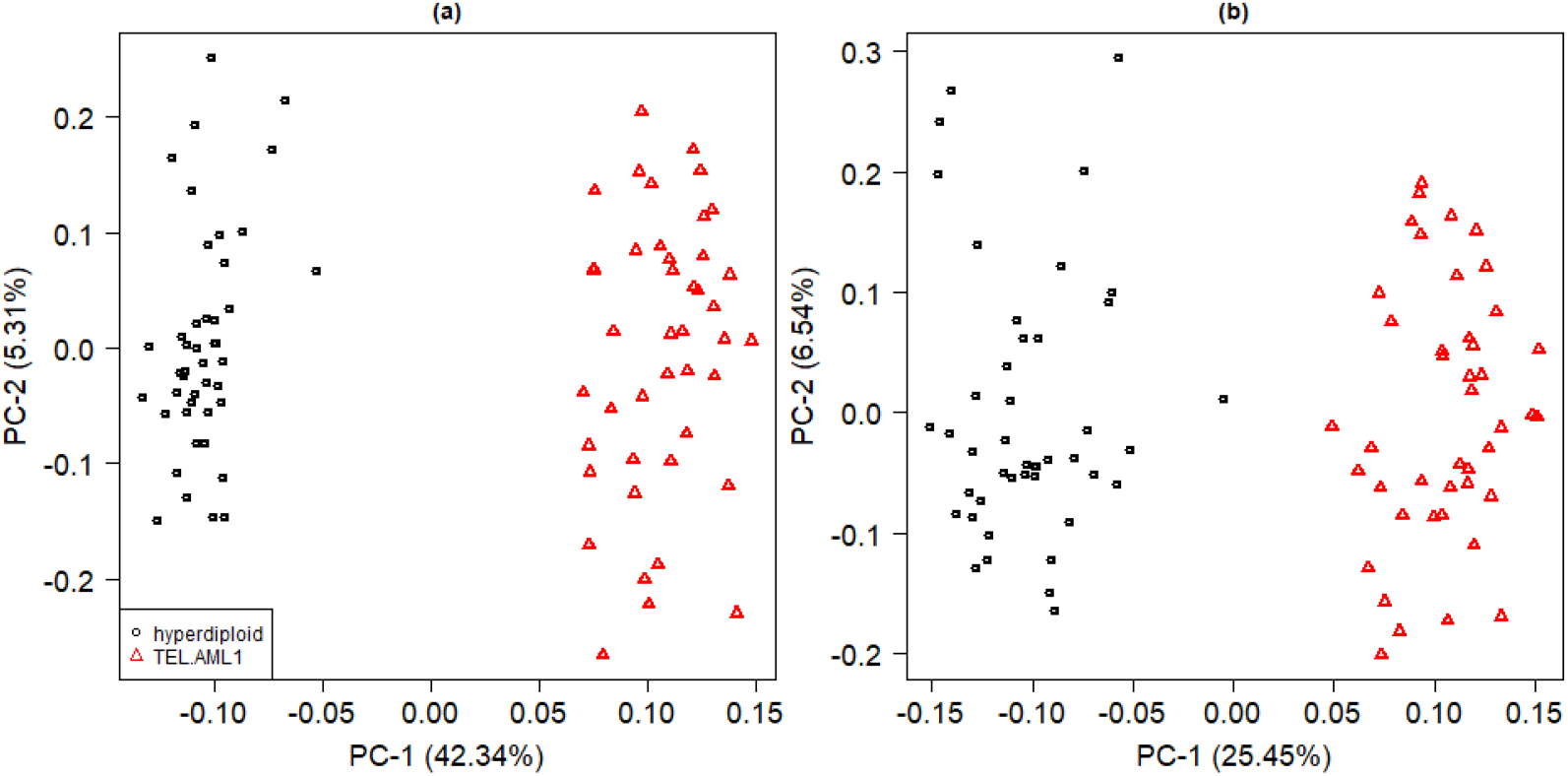
PCA plots based on the identified differentially expressed genes between hyperdiploid and TEL.AML1 subtypes of the CALL dataset using: (a) VSPCA and (b) MTT.

We again performed the analysis to evaluate the stability of the chosen superset of predictors where we reduce the number of samples in the test set of data. We used hyperdiploid samples (*n*_1_ = 44) as a training set of data and re-sampled the 43 TEL.AML1 subtype samples. The value of *n*_2_ was reduced gradually from 43 to 3 by successively subtracting 5. For all values of *n*_2_ < 43 that are considered, we selected a sample without replacement from the test set of data and conducted variable selection based on both the VSPCA and the MTT. The experiment was repeated 1000 times for each value of *n*_2_ < 43. The variables that turned out significant (at 0.01 level of significance) under smaller sample sizes that matched the ones that were selected under the entire available sample size (VSPCA selected 487 genes and MTT selected 2,196 genes as above) for the test data were counted. We display in Fig 16 the proportions of significant variables for the original complete sample with *n*_2_ = 43 that were also significant under a smaller value of *n*_2_ in more than 50% of the replicates. Again, the number of selected variables dropped much more quickly for MTT than it did for VSPCA as we reduced *n*_2_. This highlights the stability of VSPCA as well as the usefulness of VSPCA under a smaller sample size of the test set of data.

**Fig 16.**
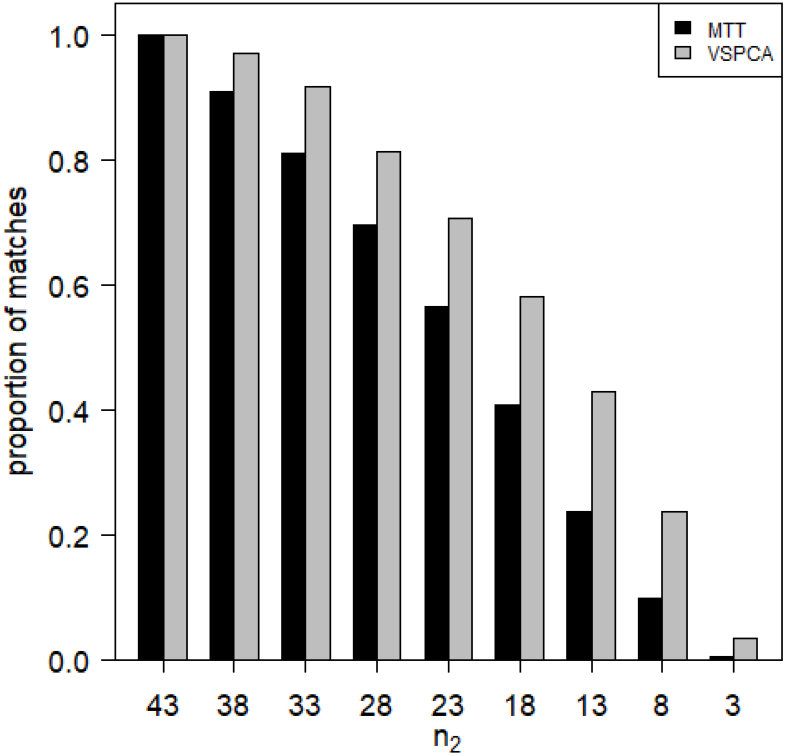
We used hyperdiploid samples (*n*_1_ = 44) as a training set of data and TEL.AML1 samples (*n*_2_ = 43) as a test set of data. The value of *n*_2_ was reduced gradually from 43 to 3 by successively subtracting 5 (as shown on the x-axis from left to right). For all values of *n*_2_ < 43 that were considered, we selected a sample without replacement from the test set of data and conducted variable selection based on VSPCA and MTT using the nominal level of significance *α* = 0.01. The experiment was repeated 1000 times for each value of *n*_2_ < 43. The variables that were significant under smaller sample sizes matched with the ones that were selected under the entire available sample size for the test data are counted. Shown are the proportions of significant variables under the entire sample size that were also significant under a smaller sample sizes in more than 50% (500/1000) of the replicates.

**Fig 17.**
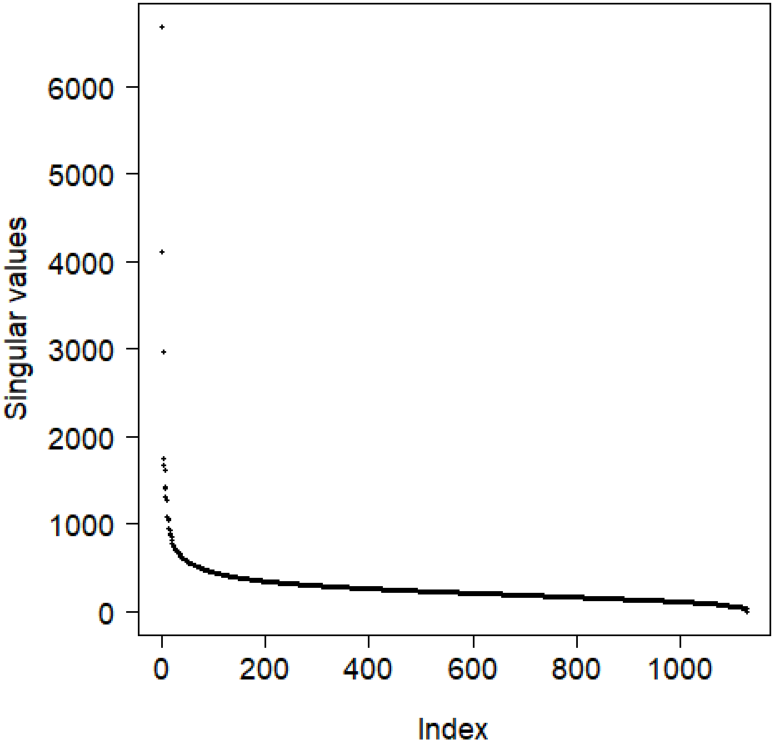
Scree plot based on 158,956 filtered markers of 1,127 accessions genotype data.

### GWAS to identify SNPs associated with rice grain length

We used a publicly available High Density Rice Array (HDRA, 700k SNPs) [46] data to further evaluate our method on high-dimensional and non-normal data. The genotype and phenotype (average grain length) data for 1,568 accessions can be downloaded from the Rice Diversity database^1^. Each of the 1,568 accessions was genotyped. The genotypic dataset of 700,000 SNPs that was assayed amounted to one SNP approximately every 0.54 kb across the rice genome. For more details about the dataset readers are referred to [46]. Note that our aim here was not to re-analyse the data but to show how well our method performs on very high-dimensional non-normal data.

We limited the genotype data to include accessions for which phenotype data are available. Out of these 1,146 accessions, we were only able to use 1,127 accessions due to the missing IDs across the phenotypes and genotypes files. These included 47 admixed, 23 admixed-indica, 102 admixed-japonica, 30 aromatic, 158 aus, 332 indica, 185 temperate-japonica, and 250 tropical-japonica sub-populations as categorized by [46]. We removed SNP loci from our analysis if the call rate was less than 95% or if minor allele frequency (MAF) is less than 5% or deviated from Hardy–Weinberg equilibrium. A total of 158,956 markers were included in the analysis none of them have more than 5% missing values. All the missing values for a SNP were set to the mean of the values pertaining to varieties whose values were non-missing for that particular SNP. The resultant genotype data, after scaling for each SNP to have zero mean and unit standard deviation, were stored in an *n* × *p* matrix *G*. We performed pre-selection PCA. The PCA plots in [46] are reproduced in Fig 18 for completeness, which shows a strong population structure. The PC-1 captured the variation between indica and japonica varieties, the PC-2 highlighted the variation between aus and indica subpopulations, and the PC-3 highlighted the variation among the three japonica subpopulations.

**Fig 18.**
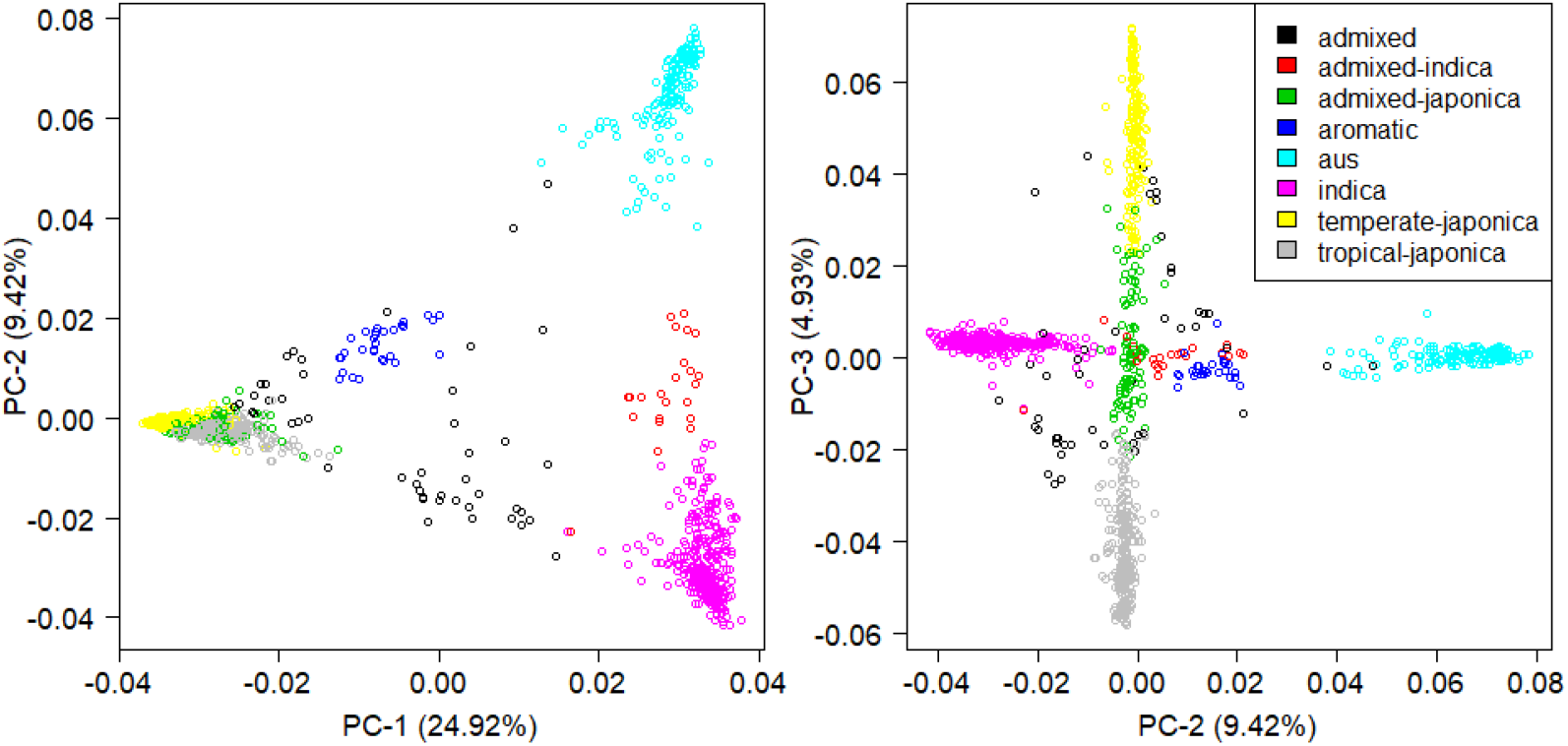
PCA plots based on the 158,956 filtered markers of 1,127 varieties of rice. The categorization into sub-populations is in accordance with [46].

The problem of confounding by population structure in genome-wide association studies (GWAS) is widely recognized [47, 26, 48]. One needs to correct for confounding factors induced by relatedness of observations to minimize FDR [47]. To correct for population stratification, we decompose *G* using a singular value decomposition and reconstruct it using *q* = 10; that is,

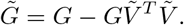

The phenotype of the 1,127 varieties had a bimodal distribution (see Fig 19) with easily distinguishable peaks and with a complete separation around the average grain length of 6 mm. Therefore, we partitioned the genotype data, contained in 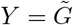, into two classes based on the phenotype variable such that the varieties of rice having average grain length of less than 6 mm fell into class-A and those having average grain length of more than 6 mm fell into class-B. We used the class-A data (*n*_1_ = 692) to obtain the estimates of the parameters. We used total variance explained criterion and set *φ* = .8. This led us to use *q* = 436 to reconstruct the genotype data from class-B (*n*_2_ = 435). The quantiles of 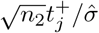 deviated from the expected quantiles under null distribution *N* (0, 1) (see Fig 21), which indicated that some SNPs are associated with the phenotype of interest. This is equally reflected in the histogram of unadjusted p-values, where the p-values are uniformly distributed except for a spike near zero (see Fig 22a).

**Fig 19.**
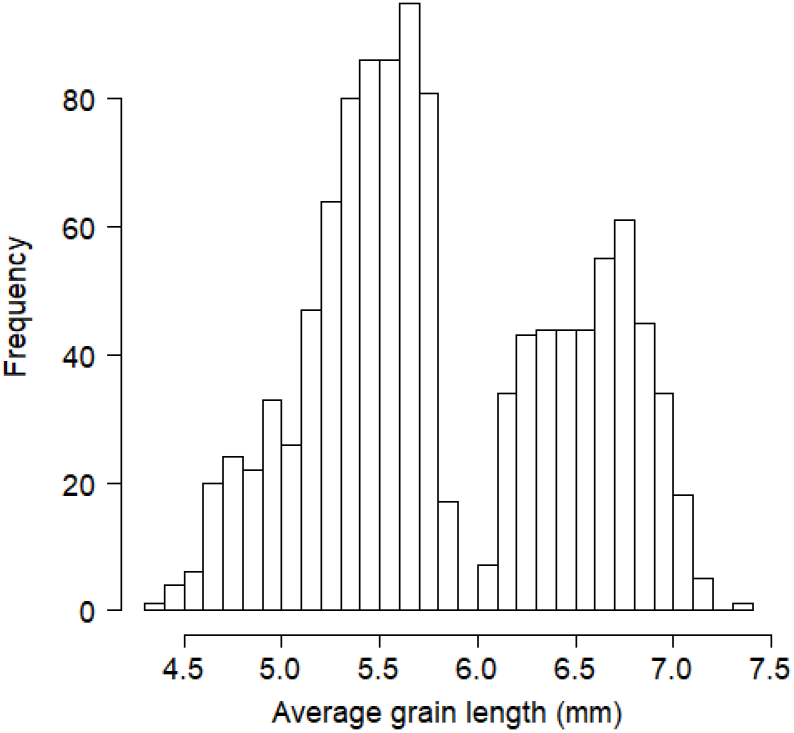
Histogram of average grain length for 1,127 varieties of rice.

**Fig 20.**
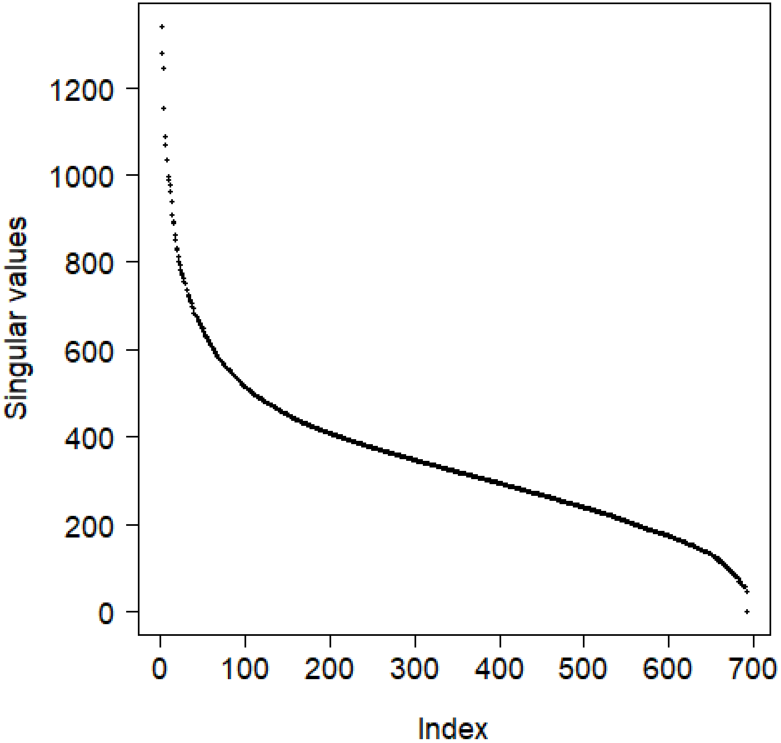
Scree plot based on 158,956 filtered markers of class-A genotype data.

**Fig 21.**
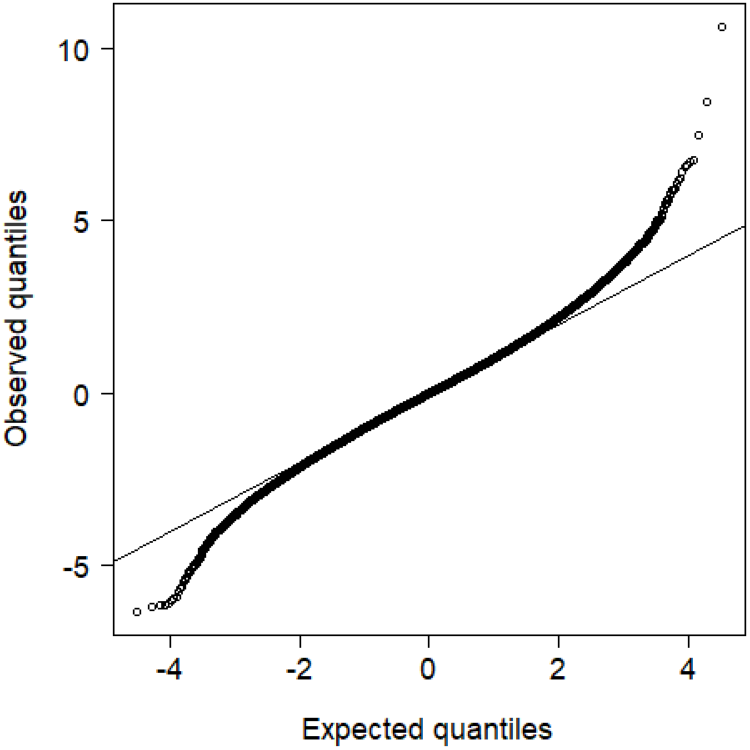
Quantiles of 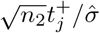 against the quantiles of standard normal distribution.

**Fig 22.**
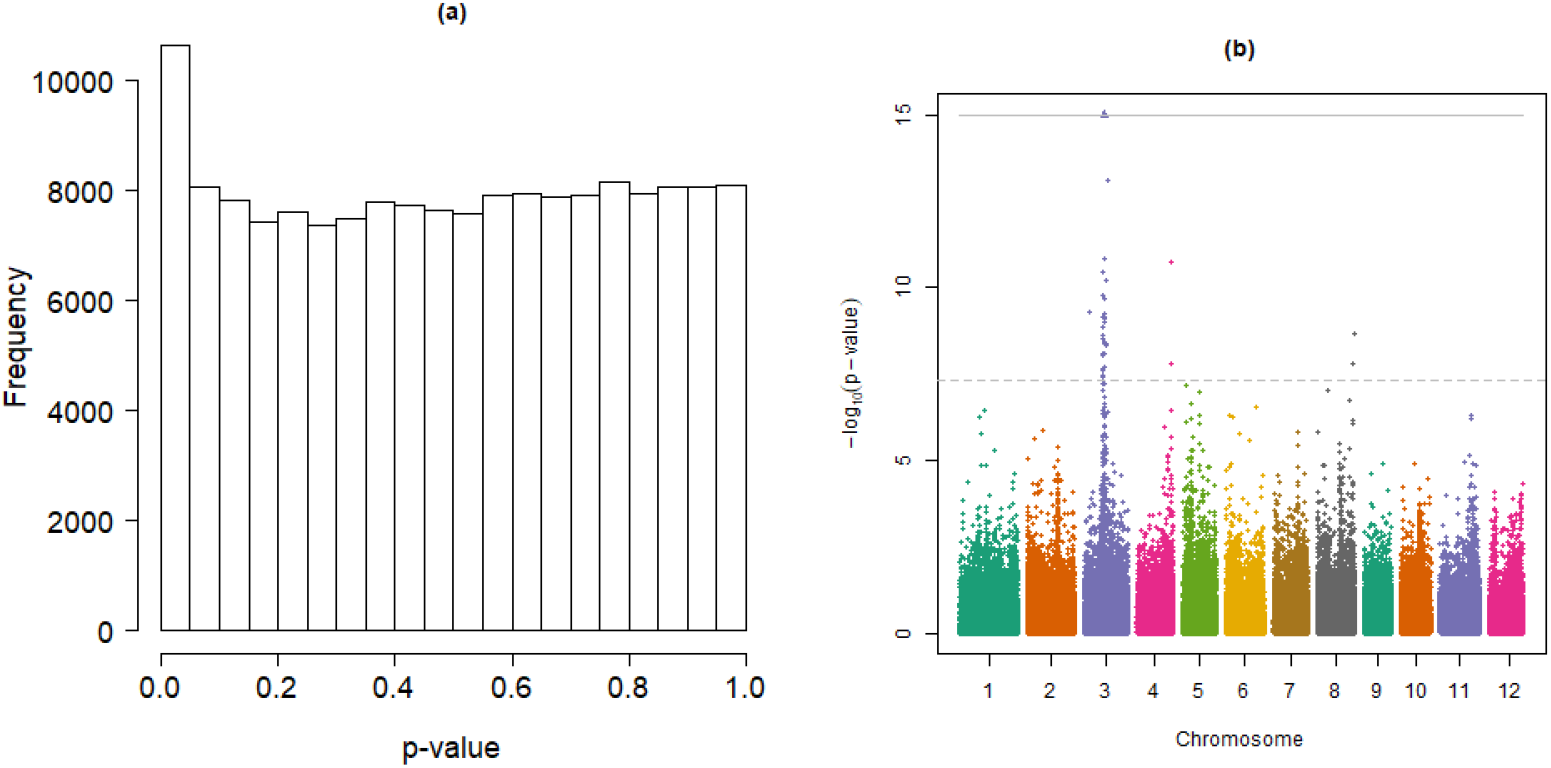
Plots of p-values: (a) Histogram of unadjusted p-values and (b) Manhattan plot based on unadjusted p-values obtained using VSPCA. The gray horizontal dashed-line indicates genome-wide significant threshold corresponding to p-value of 5*e* − 08 and the gray solid-line indicates the upper-limit of the plot along y-axis. The log_10_(p-value) for some SNPs are beyond this upper-limit and are indicated by solid-triangles.

Our GWAS identified one major locus on the rice chromosome-3 associated with the grain length and several minor loci (see Manhattan plot in Fig 22b). These findings were consistent with the previous findings. For example, the SNP-3.16732086, a functional SNP in the GS3 gene on the rice chromosome-3 [for example, see 49, 50], was found to be highly associated with grain length in [46] and is also highly significant (p-value = 2.47*e* − 26) in our analysis together with some more significant SNPs in the close vicinity on chromosome-3. The locus corresponding to the peak on chromosome-5 was found to be associated with grain size [51] and is also mentioned in [46] for its possible association with grain length. Other well pronounced peaks were located on chromosome-4 and chromosome-8, which were also observed in [46]. One reason for identifying loci on almost every chromosome could be that we have dichotomised the phenotype variable while it is used as a continuous variable in other studies [for example, see 46]. Dichotomisation of average length of grain might have confounded the association of other traits of grain with the grain length. Several quantitative trait loci (QTLs) for grain traits have been previously identified located all over the rice genome. [For an overview, see 52, 53].

## Conclusion and discussion

PCA is a classical statistical method traditionally used for dimension reduction. It is computationally manageable even for a very large dataset. A drawback of PCA, however, is that it tries to uncover the underlying structure in the data in an unsupervised manner. We overcome this drawback of PCA by taking into account the target variable. We propose to use PCA in a supervised manner for variable selection in high-dimensional datasets when the objective is class prediction in a binary classification problem such as a case-control study.

A training dataset that comes from the control group (or a group with more samples) is used to first construct a PCA model. The test dataset obtained from the cases group (or a group with a smaller sample size) is reconstructed using the low-rank approximation based on the estimates of the parameters obtained during the training phase. The induced error obtained via a low-rank reconstruction of the test set of data is used to identify and rank the variables according to their relevance to the target of interest. All variables that have the ability to predict class labels accurately will have a larger average reconstruction error and are the most informative predictors. Often we are interested in p-values associated with each variable since they are readily interpretable. Assuming that the errors are normally distributed with mean zero and with an unknown variance *σ*^2^ that we estimate using a robust estimator, we propose to obtain p-values associated with each variable. These p-values, once corrected for multiple testing, are then used for the selection of a superset of variables.

We tested our proposed method through extensive simulations, through the analysis of a well-known semi-real gene expression dataset with known differentially expressed genes and through the analysis of a real childhood acute lymphoblastic leukemia gene expression dataset. The results from a state-of-the-art independent filtering test (MTT) are also presented for comparison purposes. The proposed method identifies the superset of variables that have the ability to discriminate the classes. In contrast, the MTT is a filtering approach that evaluates feature relevance independently for each feature. Our analysis showed that the MTT is prone to choosing many redundant feature as is criticised in [2]. The method is further evaluated through the analysis of a genome-wide association sudy of a publicly available High Density Rice Array dataset. The results were consistent with the previous findings where computationally intensive mixed models were used to identify the trait loci.

PCA seeks to capture the important correlation patterns among a set of variables in the first few components. The rest of the components are believed to stand for the peculiarities and unusual observations in the data; therefore, it has been used as a core method to de-noise high-dimensional datasets. We expect our VSPCA to reduce FDR through accounting for the correlation between variables and through de-noising data by dropping a number of minor components in the training data. Disregarding the minor components will reduce the individual specific influence on the effect size since some of them are believed to represent outliers in the data.

The VSPCA is simple and computationally efficient for a high-dimensional dataset. For computational time and memory requirements of the R implementation of SVD the readers are referred to [54]. We even reduce these requirements by performing SVD only on the training data. Modern variable selection approaches such as LASSO regression require tuning parameters and computationally cumbersome methods such as cross-validation are used to tune these parameters. In contrast, our approach needs only a few lines of R code, as given in the Appendix A, and unlike LASSO regression it does not rely on tuning parameters that need to be chosen using techniques such as cross-validation, which may not be justifiable for ‘large *p* small *n*’ data. However, VSPCA requires one to choose the number of components, *q*. (indeed one can use *q* = *r*; however, we recommend dropping some minor components). We recommend choosing a value of *φ* ranging from 0.7 to 0.9 under small *n*_1_, which is a popular heuristic approach to choosing *q*. This criterion usually favours larger values of *q* and is safer when *n*_1_ is small. In our experience, changing the value of *φ* from 0.7 to 0.9 did not make a noticeable difference to the final results. Other heuristics such as looking for an elbow point in a scree plot could be used as well. However, we found that the scree plot is less safe because it favours smaller values of *q* and an extra-small value of *q* might have a large effect on the results. We found it safe to use a slightly larger value of *q* (compared to what the elbow point suggest) since it does not make a notable change to the results. Appropriate choices of *q* for PCA in general, and for VSPCA in particular, is also worthy of future investigation.

## Supporting information

**Supplementary materials: Full set of simulation results.**

## Appendix: A R code to perform simulations

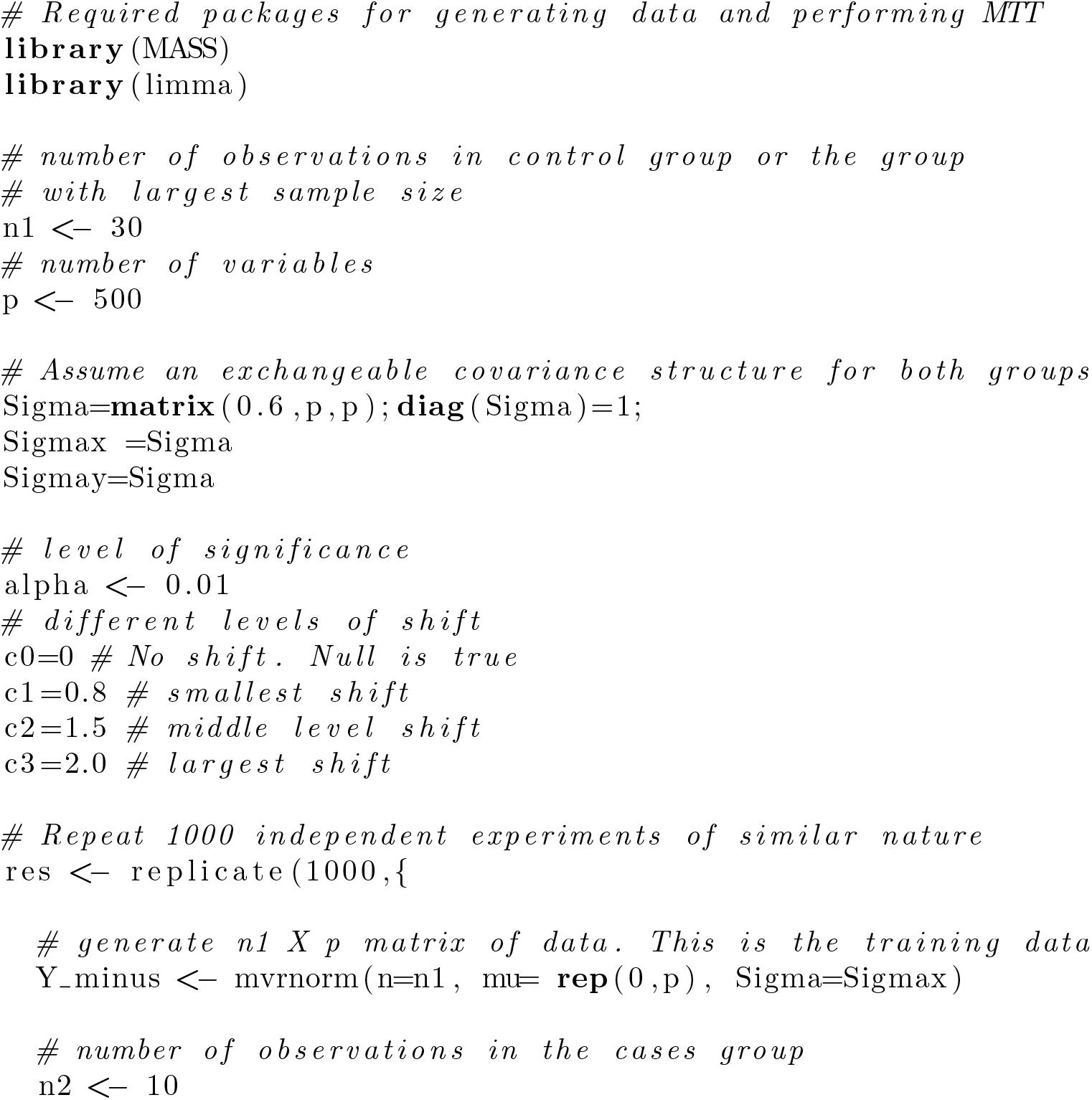

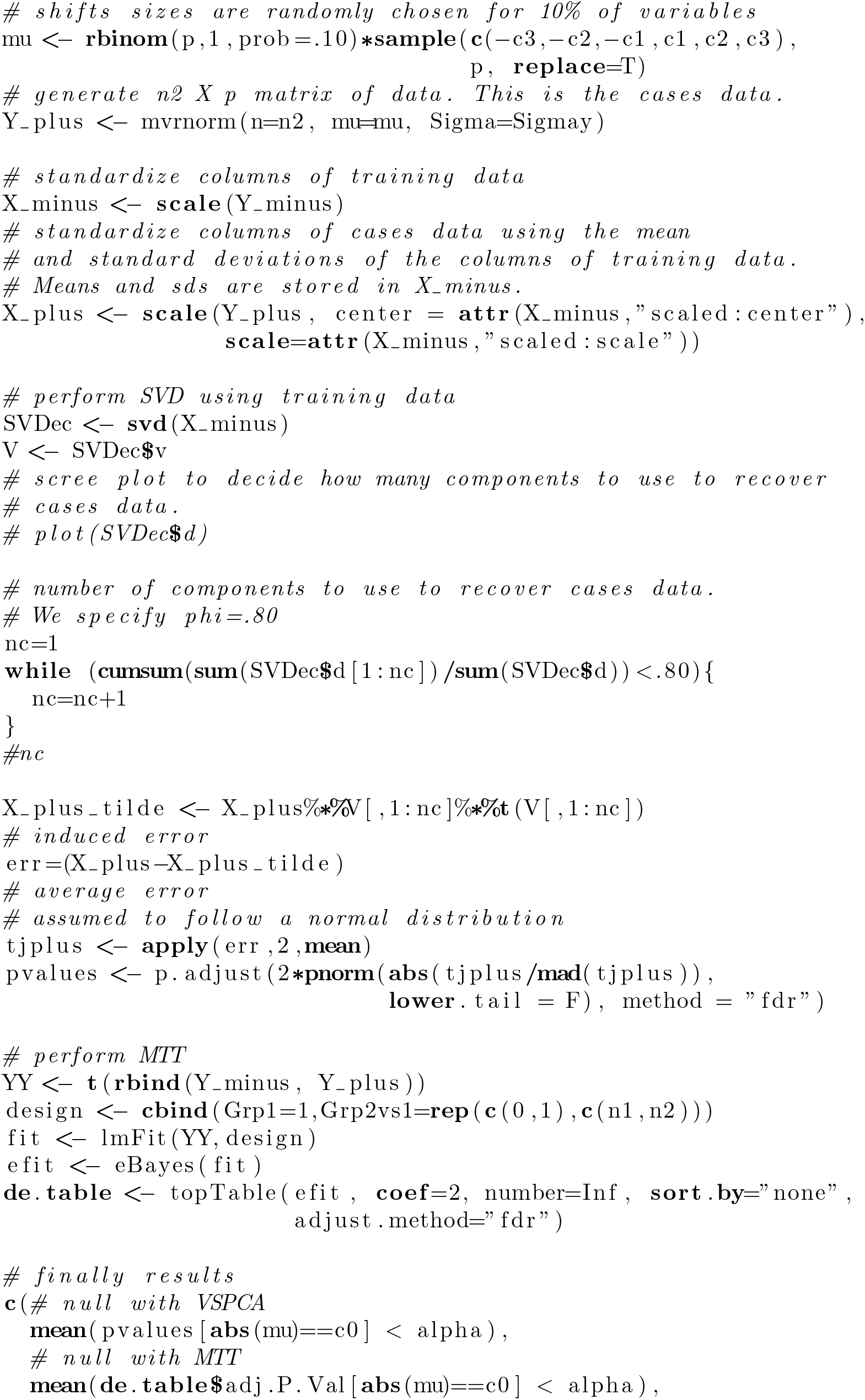

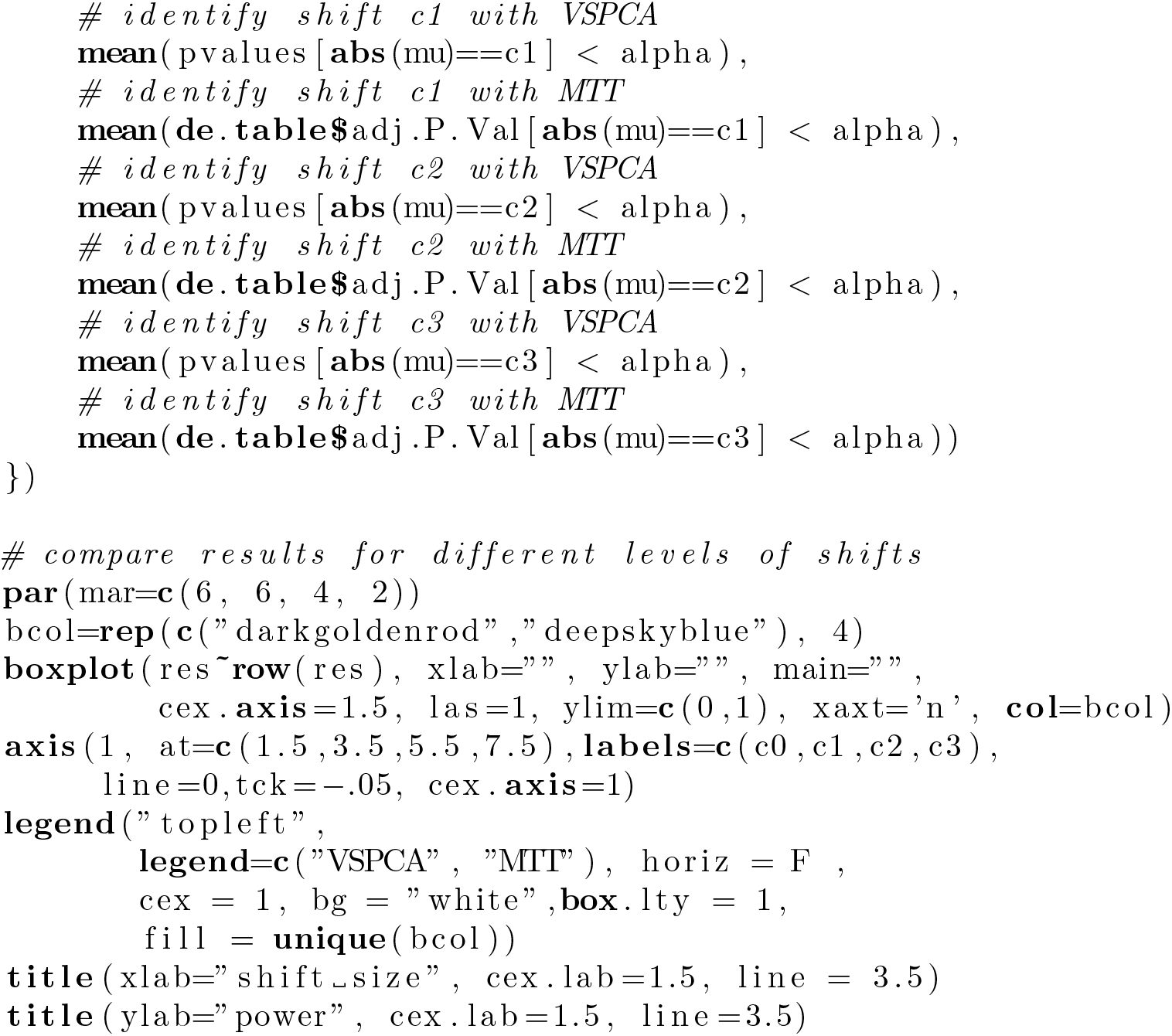

## Acknowledgments

This research was supported by an ARC Australian Laureate Fellowship under Grant No. FL150100150. The authors also acknowledge the support of the Australian Research Council (ARC) Centre of Excellence for Mathematical and Statistical Frontiers (ACEMS).

## Footnotes

1 http://www.ricediversity.org/

